# Synaptotagmin 1 mediates toxicity of botulinum neurotoxin type A

**DOI:** 10.1101/2022.02.28.482398

**Authors:** Merja Joensuu, Vanessa Lanoue, Parnayan Syed, Tristan P. Wallis, James Rae, Ailisa Blum, Rachel Gormal, Christopher Small, Shanley Sanders, Anmin Jiang, Stefan Mahrhold, Nadja Krez, Michael Cousin, Ruby Cooper-White, Justin J. Cooper-White, Brett M. Collins, Robert G. Parton, Giuseppe Balistreri, Andreas Rummel, Frédéric A. Meunier

## Abstract

The extreme neurotoxicity of botulinum neurotoxins (BoNTs), broadly used as a therapeutics, is mediated by their direct binding to two plasma membrane receptors: polysialogangliosides (PSGs) and serotype-dependently either synaptotagmin 1/2 (Syt1/2) or synaptic vesicle glycoprotein 2 (SV2). Here, we demonstrate that although BoNT/A serotype binds directly to PSG and SV2, its neurotoxicity depends on Syt1. Using single-molecule superresolution microscopy of pathological concentrations of BoNT/A holotoxins, we demonstrate that the toxin binds to a preassembled PSG-Syt1 complex that forms nanoclusters with SV2 on the plasma membrane of hippocampal neurons. This coincidental interaction controls the selective endocytic targeting of BoNT/A into synaptic vesicles, leading to incapacitation of neuronal communication. Our results suggest that Syt1-SV2 plasma membrane nanoclusters may acts as a common target for various BoNT serotypes.

**One-Sentence Summary:** BoNT/A hijacks an intrinsic tripartite PSG-Syt1-SV2 endocytic sorting mechanism to incapacitate neuronal communication

---

The extreme potency of clostridial neurotoxins (CNTs), including botulinum neurotoxins (BoNTs), is mediated by their high specificity to peripheral cholinergic motorneurons, leading to flaccid paralysis and respiratory failure (*1*). Targeted entry of BoNTs into synaptic vesicles (SVs) is critical for their toxic function. All BoNTs share a similar molecular structure consisting of a heavy chain (HC), connected to a light chain (LC) by a disulfide bond and non-covalent interactions (*2*). The HC drives the specific uptake of BoNTs and, following the acidification of the SV lumen, promotes the LC translocation into the cytosol where it cleaves its target soluble N-ethylmaleimide sensitive factor attachment protein receptor (SNARE) thereby blocking neurotransmission (*3*). The dual receptor model stipulates that BoNTs first bind to the abundant plasma membrane polysialogangliosides (PSGs) with low affinity, and then with high affinity to a specific protein receptor (*4*). BoNT/A and E bind to N-glycosylated SV glycoprotein 2 (SV2), while BoNT/B, DC, and G bind to synaptotagmin 1/2 (Syt1/2) (*5*), as a requisite step for endocytosis into SVs. Plasma membrane Syt1-SV2 nanoclusters control SV recycling (*6–8*), suggesting that they may act as a common target of BoNTs for selective targeting into SVs.

To determine if BoNT/A uses Syt1-SV2 nanoclusters to internalize into neurons, we first investigated the effects of Syt1 knockdown (KD) on the neurotoxic function of BoNT/A. Using immunofluorescence assays and a lentiviral CRISPR interference (CRISPRi) tool with short guide RNAs (sgRNAs) 1-3 (Fig. 1A-C and fig. S1A), and short hairpin RNA (shRNA) in cultured hippocampal neurons (fig. S1B-H), we found that Syt1 KD was protective against Abobotulinumtoxin A (a BoNT/A-based licensed drug)-induced cleavage of the SNARE protein synaptosomal-associated protein 25kDa (SNAP-25), a hallmark of BoNT/A-induced intoxication (*3*). Syt1 re-expression in the Syt1 sgRNA1 KD background rescued the phenotype (Fig. 1D-F), and Syt1 overexpression in wildtype neurons led to a significant increase in SNAP-25 cleavage following toxin treatment (Fig. 1H-J), indicating a gain of toxic function and suggesting that the expression level of Syt1 may be rate limiting for BoNT/A intoxication. We validated the protective effect of Syt1 sgRNA1 KD on BoNT/A function by whole-cell patch-clamp recordings of spontaneous miniature postsynaptic currents (mIPSCs) in hippocampal neurons (Fig. 1K). Abobotulinumtoxin A treatment did not affect mIPSC amplitudes across the conditions (Fig. 1L) and Syt1 sgRNA1 KD was protective over the mIPSCs frequency reduction observed in control neurons (Fig. 1M), demonstrating that Syt1 sgRNA1 KD leads to loss of BoNT/A-induced toxicity.

**Fig. 1.**
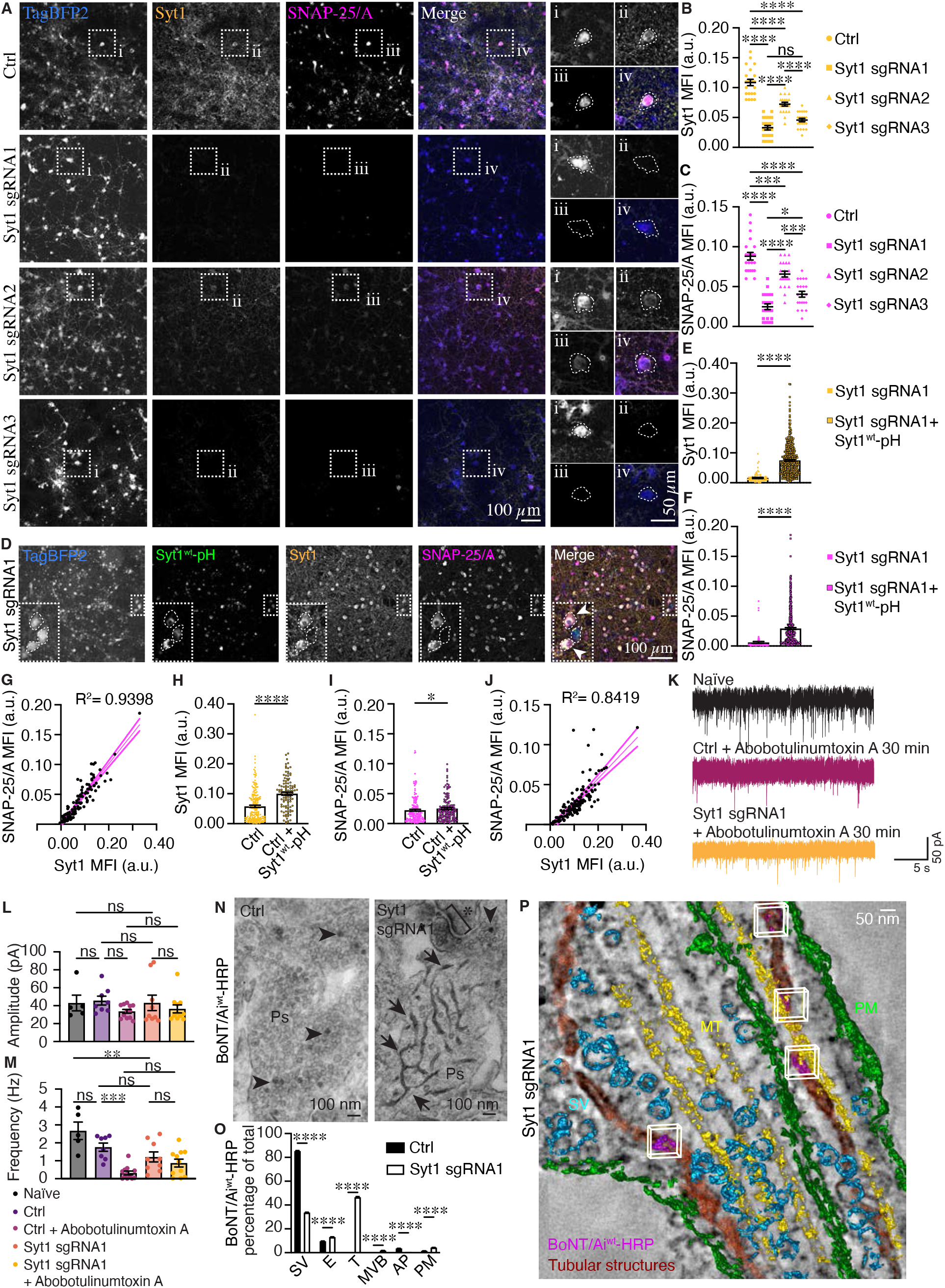
CRISPRi knockdown of Syt1 leads to deficient endocytic uptake of BoNT/A into synaptic vesicles and loss of BoNT/A toxic function. (**A**) Confocal Z-stack sum projections of neurons transduced with TagBFP2-expressing (blue) control (ctrl; non-targeting sgRNA) and CRISPRi Syt1 KD (sgRNA1-3) lentiviruses. Neurons were treated for 30 min with 10 units of Abobotulinumtoxin A, and immunostained for endogenous Syt1 (yellow) and BoNT/A-cleaved SNAP-25 (SNAP-25/A; magenta). Boxed regions (i-iv) are magnified on right. Neuronal somas are outlined with dashed lines for clarity. Quantification of mean fluorescence intensity (MFI) of endogenous (**B**) Syt1 and (**C**) SNAP-25/A in control and Syt1 sgRNA1 KD neurons following 30 min Abobotulinumtoxin A (10 units) treatment. (**D**) Confocal Z-stack sum projection of neurons transduced with TagBFP2-expressing (blue) CRISPRi Syt1 KD (sgRNA1) lentiviruses and rescued with pLenti6.3-Syt1^wt^-pH (green). Neurons were treated with Abobotulinumtoxin A (10 units) for 30 min, and immunostained for endogenous Syt1 (yellow) and SNAP-25/A (magenta). Boxed regions are magnified in the bottom left corners; rescued neurons are indicated with arrowheads and neuronal somas are outlined with dashed lines for clarity. Quantification of MFI of endogenous (**E**) Syt1 and (**F**) SNAP-25/A in Syt1 sgRNA1 KD neurons, with and without pLenti6.3-Syt1^wt^-pH rescue, and treated with Abobotulinumtoxin A (10 units) for 30 min. (**G**) Linear regression with 95% confidence intervals (dotted lines) of Syt1 and SNAP-25/A MFI is shown from a representative acquisition. Quantification of MFI of endogenous (**H**) Syt1 and (**I**) SNAP-25/A in non-transduced neurons (neurons negative for TagBFP2) with and without pLenti6.3-Syt1^wt^-pH expression following Abobotulinumtoxin A treatment (10 units) for 30 min. (**J**) Linear regression with 95% confidence intervals (dotted lines) of Syt1 and cleaved SNAP-25/A MFI is shown from a representative acquisition. (**K**) Representative patch-clamp spontaneous miniature postsynaptic current (mIPSC) recordings in naïve neurons, and in control (non-targeting) and Syt1 KD (CRISPRi sgRNA1) neurons following 30 min treatment with Abobotulinumtoxin A (10 units). Quantification of mIPSC (**L**) peak amplitudes and (**M**) frequencies for indicated conditions. (**N**) EM analysis of endocytosed BoNT/Ai^wt^-HRP following lentiviral control (ctrl; non-targeting sgRNA) and Syt1 sgRNA1 KD transduction. HRP precipitate in synaptic vesicles (SVs; arrowheads) and tubular structures (arrows) in presynapses (Ps) are indicated. The postsynaptic density is marked with an asterisk. (**O**) Percentage of total BoNT/Ai^wt^-HRP localized in SVs, endosomes (E), tubular structures (T), multivesicular bodies (MVB), structures resembling autophagosomes (AP) and on the plasma membrane (PM) in indicated conditions. (**P**) Electron tomogram of endocytosed BoNT/Ai^wt^-HRP (purple with white bounding boxes) in tubular structures (red) induced by Syt1 sgRNA1 KD. PM (green), SVs (cyan), microtubules (MT; yellow). (A-C) N=21 acquisitions/condition from 2 independent experiments, (E,F,H,I) n=3 acquisitions/condition from 1 experiment, (K,L) n=5-12 acquisitions/condition from 2 independent experiments and (M,N) n=56 ROIs/condition from 2 independent experiments. Non-parametric Kruskal-Wallis multiple comparison test (B,C,L), ordinary one-way ANOVA multiple comparison test (M) and non-parametric Mann-Whitney U test (E,F,H,I,O). *p<0.05, **p<0.01, ***p<0.001, ****p<0.0001 and n.s, non-significant.

Having obtained evidence that Syt1 mediates BoNT/A toxicity, we examined the endocytic uptake of BoNT/A following Syt1 KD using electron microscopy (EM) analysis and catalytically inactivated (E224A/R363A/Y366F) wildtype (BoNT/Ai^wt^) holotoxin (*9*) (fig. S2A and Movie S2) labeled with horseradish peroxidase (HRP) for cytochemical staining. Following activity-dependent uptake of BoNT/Ai^wt^-HRP in cultured hippocampal neurons, the majority of the toxin was internalized into SVs in control neurons, whereas in Syt1 sgRNA1 KD neurons, BoNT/Ai^wt^-HRP was found in tubular structures (Fig. 1M,N). These tubular structures, which were confirmed using electron tomography (Fig. 1P and Movie S1), occasionally originated from the plasma membrane, where they remained open to the extracellular space (fig. S1I). Similar tubulation was not observed following Syt1 sgRNA1 KD without BoNT/Ai exposure (Ruthenium red staining; fig. S1J), indicating either that the tubules were induced specifically by the toxin, or that the toxin was internalized into tubular endosomes that had pinched off from the plasma membrane. Together these results suggest that the neurotoxic protection afforded by Syt1 KD is due to the accumulation of BoNT/A in tubules which presumably lack the acidic environment (*10*) that is required for the BoNT/A-LC to translocate into the cytosol and cleave SNAP-25 (*11*).

Membrane receptors exhibit lateral diffusion within the plasma membrane (*12, 13*), and can agglomerate into nanoclusters, with restricted diffusion, that have a functional role in ligand binding (*14*). To provide evidence that the direct binding of BoNT/A to SV2 affects their nanoscale organization and mobility on the plasma membrane, we first performed dual-color single-molecule imaging using universal point accumulation imaging in nanoscale topography (uPAINT) technique (*15*). Hippocampal neurons transiently expressing pHluorin-tagged SV2A^wt^ (*16*) (SV2A^wt^-pH) were stimulated with high K^+^ buffer supplemented with 100 pM anti-GFP Atto565 nanobodies (At565nb) and 100 pM BoNT/Ai^wt^ labeled with Atto647N (BoNT/Ai^wt^-At647N; fig. S2A and Movie S2) which, based on e*x vivo* mouse phrenic nerve assays, did not prevent the BoNT/A neurotoxicity (fig. S2B-S2E, fig. S3A,S3E). The mobility of SV2A^wt^-pH-bound At565nb was then recorded simultaneously with the toxin (Movie S3), and the two were occasionally observed confined in the same regions of interest (ROIs), exhibiting restricted lateral diffusion (Fig. 2A) which, based on Nanoscale Spatiotemporal Indexing Clustering (NASTIC) analysis (*17*), reflected co-clustering of SV2A^wt^-pH/At565nb and BoNT/Ai^wt^-At647N (Fig. 2B). Prior to co-clustering, both the toxin and SV2A were mobile on the plasma membrane with distinct mobility (the mobility of the toxin being higher than that of its receptor), and following lateral trapping in nanoclusters, the mobility of both decreased and became indistinguishable from one another (Fig. 2B), suggesting that SV2 is a dynamic, rather than static, target of the toxin. In support, SV2A^wt^-pH/At565nb single-molecule mobility decreased significantly following BoNT/Ai^wt^-At467N treatment (Fig. 2C-2E, fig. S4A-S4C), which was not observed with a BoNT/Ai mutant with abolished binding to SV2 (BoNT/Ai^SV2^-At647N; G1141D/G1292R mutations (*18, 19*)) (Fig. 2C, fig. S2A, fig. S3C,S3E, fig. S4D-H, and Movie S2) demonstrating that the effect on receptor mobility was directly induced by BoNT/A. Introducing the SV2A^T84A^ mutation, that prevents Syt1-SV2 interactions (*20*), greatly reduced this effect (SV2A^T84A^-pH/At565nb; Fig. 2C, fig. S4I-S4M), indicating that the interaction between Syt1 and SV2 plays a role in the toxin-induced confinement of SV2A. In support, we observed a similar change in the mobility of Syt1^wt^-pH/At656nb (Syt1^wt^-pHluorin(*21*)) following BoNT/Ai^wt^-At647N exposure (Fig. 2F and fig. S5A-S5D), which was surprising given that Syt1 does not directly bind to BoNT/A (*22*), indicating that the toxin-induced effects on Syt1 mobility could be indirect. Syt1 was recently shown to form a complex with GT1b ganglioside (*23*). This complex, that functions as a receptor for BoNT/B, was disrupted with Syt1^K52A^ mutation, leading to deficient binding and entry of BoNT/B (*23*). Given that BoNT/B shares a conserved PSG-binding site with BoNT/A (*24*), we hypothesized that a similarly preassembled PSG-Syt1 complex may also be hijacked by BoNT/A to mediate the observed confinement of Syt1. Indeed, BoNT/Ai^wt^-At647N treatment did not change Syt1^K52A^-pH/At565nb uPAINT mobility (Fig. 2F. fig. S5E-S5H), indicating that the toxin-induced effects on the plasma membrane mobility of Syt1 occurred via binding to the ganglioside in the GT1b-Syt1 complex. To address whether the interactions between Syt1 and SV2 contributed to the BoNT/A-induced confinement of Syt1, we repeated the uPAINT experiments with Syt1^K326A,K328A^-pH/At565nb, which is a mutant that prevents Syt1 binding to SV2 (*16*), and observed a markedly reduced mobility of Syt1 (Fig. 2D and fig. S5I-S5L). In support of these findings, BoNT/Ai^wt^-At647N treatment increased the SV2A^wt^-pH/At565nb nanoclustering (Fig. 2G-I), and decreased the density of tracks within a cluster as well as the cluster size (fig. S6A,6B). Similar effects were not observed with SV2A^T84A^-pH/At565nb (Fig. 2J and fig. S6C,S6D). BoNT/Ai^wt^-At647N treatment also affected Syt1^wt^-pH/At565nb nanoclustering (Fig. 2K and fig. S6I-S6N), but not that of Syt1^K52A^-pH/At565nb (Fig. 2L and fig. S6I,S6J) or Syt1^K326A,K328A^-pH/At565nb (Fig. 6M and fig. S6K,S6L). The increased SV2A and Syt1 nanoclustering on the plasma membrane correlated with increased Syt1-SV2 protein-protein interactions that we measured using the *in situ* proximity ligation assay (PLA; Fig. 2N-O). First we generated a W1266L (*25*) mutation in BoNT/Ai to abolish toxin binding to PSG (BoNT/Ai^PSG^), as well as an additional double mutant with abolished binding to both PSG and SV2 (G1141D/W1266L/G1292R; BoNT/Ai^PSG,SV2^) (fig. S2A and Movie S2), and then labeled the mutant holotoxins with At647N (fig. S3C-S3E). The discrete fluorescent PLA dots provides a readout of a productive ligation of endogenous Syt1 and SV2C isoform, when proteins are in <40 nm proximity to one another. Treatment of hippocampal neurons with BoNT/Ai^wt^-At647N significantly increased the number of Syt1-SV2C PLA dots in the somato-dendritic area of the neurons compared to naïve or high K^+^-stimulated neurons, or following exposure to BoNT/Ai^PSG^-At647N, BoNT/Ai^SV2^-At647N or BoNT/Ai^PSG,SV2^-At647N diluted in high K^+^ (Fig. 2N-O). Together, these results indicate that the intrinsic PSG-Syt1-SV2 interactions are a requisite for BoNT/A-induced receptor confinement and nanoclustering, and that these effects are induced only when BoNT/A coincidentally binds to PSG and SV2.

**Fig. 2.**
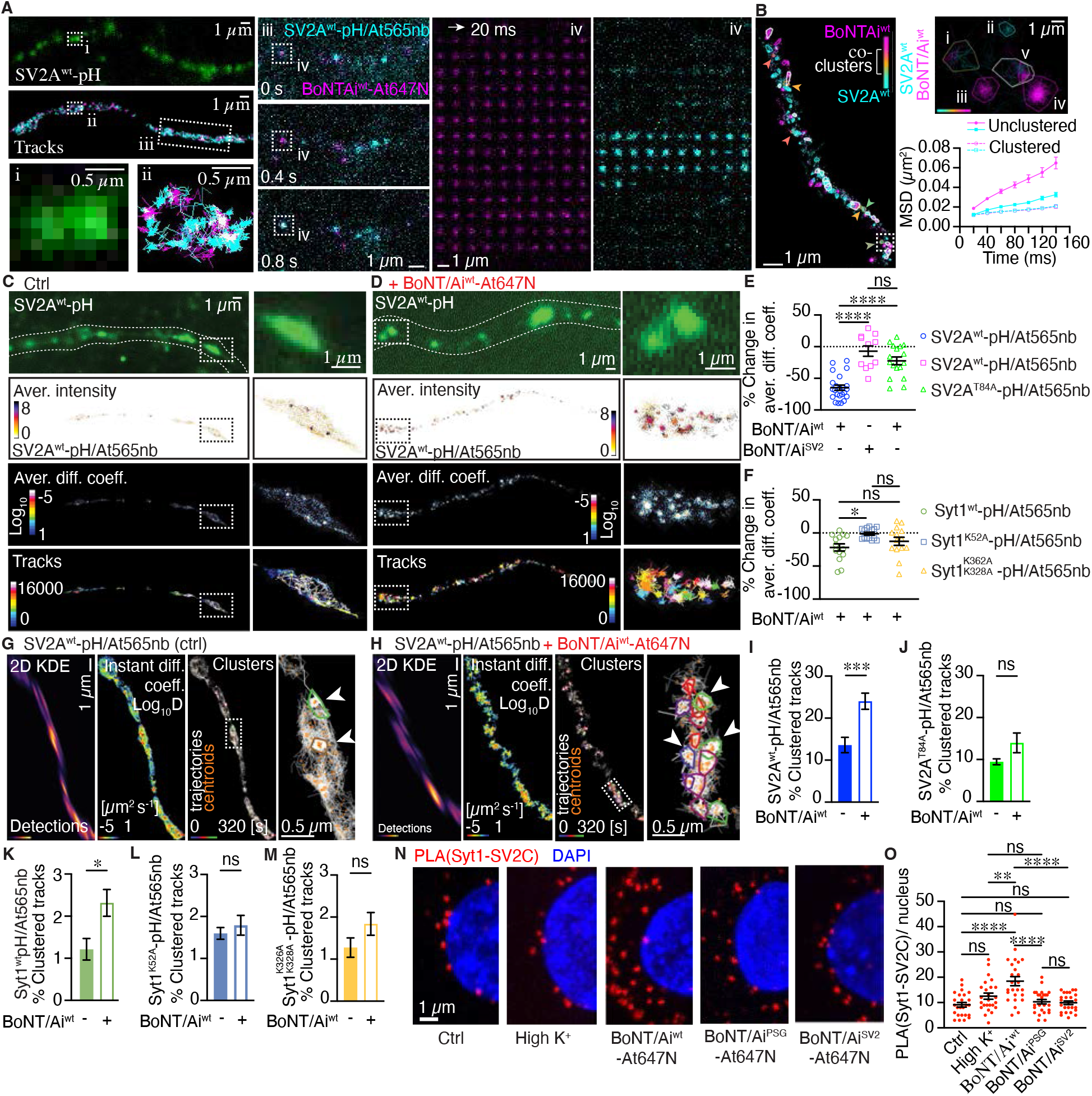
BoNT/A plasma membrane binding induces SV2 and Syt1 receptor clustering. (**A**) Dual color super-resolution imaging of a live hippocampal neuron expressing SV2A^wt^-pH (green), and corresponding uPAINT trajectories of SV2A^wt^-pH/At565nb (cyan) imaged simultaneously with BoNT/Ai^wt^-At647N (magenta). Boxed regions (i-ii) are magnified in the bottom, and a selected time-lapse sequence of the single molecules of SV2A^wt^-pH/At565nb and BoNT/Ai^wt^-At647N detected coincidentally in a neuronal projection from region (iii) is shown as indicated. Frame-by-frame coincidental detection of BoNT/Ai^wt^-At647N and SV2A^wt^-pH/At565nb from the boxed 10 x 10 pixel region (iv) is displayed from a 50 Hz (20 ms exposure time) 16,000 frame acquisition as indicated. (**B**) Representative NASTIC cluster analysis image of BoNT/Ai^wt^-At647N and SV2A^wt^-pH/At565nb co-clustering indicated by colored arrowheads, and hotspots with repeated nanoclusters by white outlines. Boxed area is shown magnified on left, depicting (i) a BoNT/Ai^wt^-At647N and SV2A^wt^-pH/At565nb co-cluster, (ii) SV2A^wt^-pH/At565nb cluster, (iii,v) BoNT/Ai^wt^-At647N cluster, and a hotspot (v) as indicated. Mean square displacement (MSD) graph shows unclustered and clustered BoNT/Ai^wt^-At647N (magenta) and SV2A^wt^-pH/At565nb (cyan). Neurons expressing SV2A^wt^-pH were imaged with uPAINT in (**C**) control conditions (high K^+^) and (**D**) following BoNT/Ai^wt^-At647N treatment (diluted in high K^+^) showing low-resolution images of SV2A^wt^-pH (green) and super-resolved average intensity, diffusion coefficient and trajectory maps of SV2A^wt^-pH/At565nb. Boxed areas magnified on right. (**E**) Percentage change in SV2A^wt^-pH/At565nb and SV2A^T84A^-pH/At565nb uPAINT mobility following BoNT/Ai^wt^-At647N or BoNT/Ai^SV2^-At647N treatments, normalized to the average mobility of the control condition (high K^+^; non-treated). (**F**) Percentage change in Syt1^wt^-pH/At565nb, Syt1^K52A^-pH/At565nb and Syt1^K326A,K328A^-pH/At565nb mobility following BoNT/Ai^wt^-At647N treatment, normalized to the average of the control (high K^+^; non-treated). Representative images of NASTIC nanocluster analysis of SV2A^wt^-pH/At565nb in (**G**) control condition (high K^+^) and (**H**) following high K^+^ stimulation in the presence of BoNT/Ai^wt^-At647N. Boxed areas are magnified on right, arrowheads indicate nanoclusters. Nanocluster quantification of (**I**) SV2A^wt^-pH/At565nb, (**J**) SV2A^T84A^-pH/At565nb, (**K**) Syt1^wt^-pH/At565nb, (**L**) Syt1^K52A^-pH/At565nb and (**M**) Syt1^K326AA,K328A^-pH/At565nb in control conditions (high K^+^;“-”) and following BoNT/Ai^wt^-At647N treatment (+). (**N**) *In situ* proximity ligation assay (PLA) of Syt1-SV2C interactions in naïve (ctrl) and high K^+^ stimulated (vehicle) conditions, and following high K^+^ stimulation with the indicated toxins. Nucleus (DAPI, blue) and Syt1-SV2C PLA (red) signals from a representative confocal maximum intensity projections of the somatodendritic area. (**O**) Number of PLA (Syt1-SV2C) dots per nucleus in the indicated conditions. (E) N=11-23 acquisitions/condition from 3-5 independent experiments, (F) n=14 acquisitions/condition from 3-4 independent experiments, (IM) n=11-19 acquisitions/condition from 3-5 independent experiments, and (O) n=25 ROIs (200×200 pixel ROI surrounding DAPI-staining)/condition from 2 independent experiments. (C,D) Non-parametric Kruskal-Wallis multiple comparison test, (I,J,K,L) parametric unpaired *t* test, and (M) non-parametric Mann-Whitney U test. *p<0.05, **p<0.01, ***p<0.001, ****p<0.0001, and n.s, non-significant.

To determine whether coincidental binding to PSG and SV2 was a requisite for BoNT/A endocytosis, we performed single-molecule imaging of BoNT/Ai^wt^-At647N and mutants on the plasma membrane (uPAINT) and following activity-dependent endocytosis using a pulse-chase technique called subdiffractional tracking of internalized molecules (sdTIM; Fig. 3A, Movie S4) (*26*). BoNT/Ai^wt^-At647N mobility decreased significantly following internalization (Fig. 3B and fig. S7A-S7C), reflecting endocytosis into SVs (*27*), whereas similar decrease was not observed with the mutated toxins (Fig. 3B and fig. S7D-S7L), indicating defective endocytosis. Abolishing BoNT/Ai binding to PSGs significantly decreased the number of single-molecule trajectories on the plasma membrane compared to BoNT/Ai^wt^-At647N, with a similar trend also being observed with BoNT/Ai^SV2^-At647N and BoNT/Ai^PSG,SV2^-At647N (Fig. 3C), and following activitydependent uptake of the toxins (Fig. 3D). Uptake of the toxins, followed by fixation and a fluorescence emission assay, demonstrated a significant decrease in the fluorescence emission steps of all the receptor-binding mutants compared to BoNT/Ai^wt^-At467N in presynaptic areas (fig. S7M-O). To verify the internalization deficiency exerted by the mutations, we next conducted a pulse-chase uptake of HRP-labeled BoNT/Ai^wt^ and receptor binding mutants, after which the neurons were cytochemically stained and processed for EM. The great majority of the BoNT/Ai^wt^-HRP was observed internalized into SVs, with only a small portion localizing in other endocytic compartments or remaining on the plasma membrane (Fig. 3E,3F). In contrast, most of the BoNT/Ai mutants were found on the plasma membrane (Fig. 3Eii), demonstrating a defect in the endocytic uptake of the mutant toxins (Fig. 3F). The average sectional areas of the BoNT/Ai^wt^-HRP–stained endocytic compartments were significantly smaller than those of the mutants (Fig. 3G). Using a retrograde transport assay, internalized BoNT/Ai^wt^-At647N was also observed trafficking from the nerve terminals to the cell soma, while the retrograde carrier frequency of BoNT/Ai mutants was significantly decreased (Figure 3H-3J; Movie S5).

**Fig. 3.**
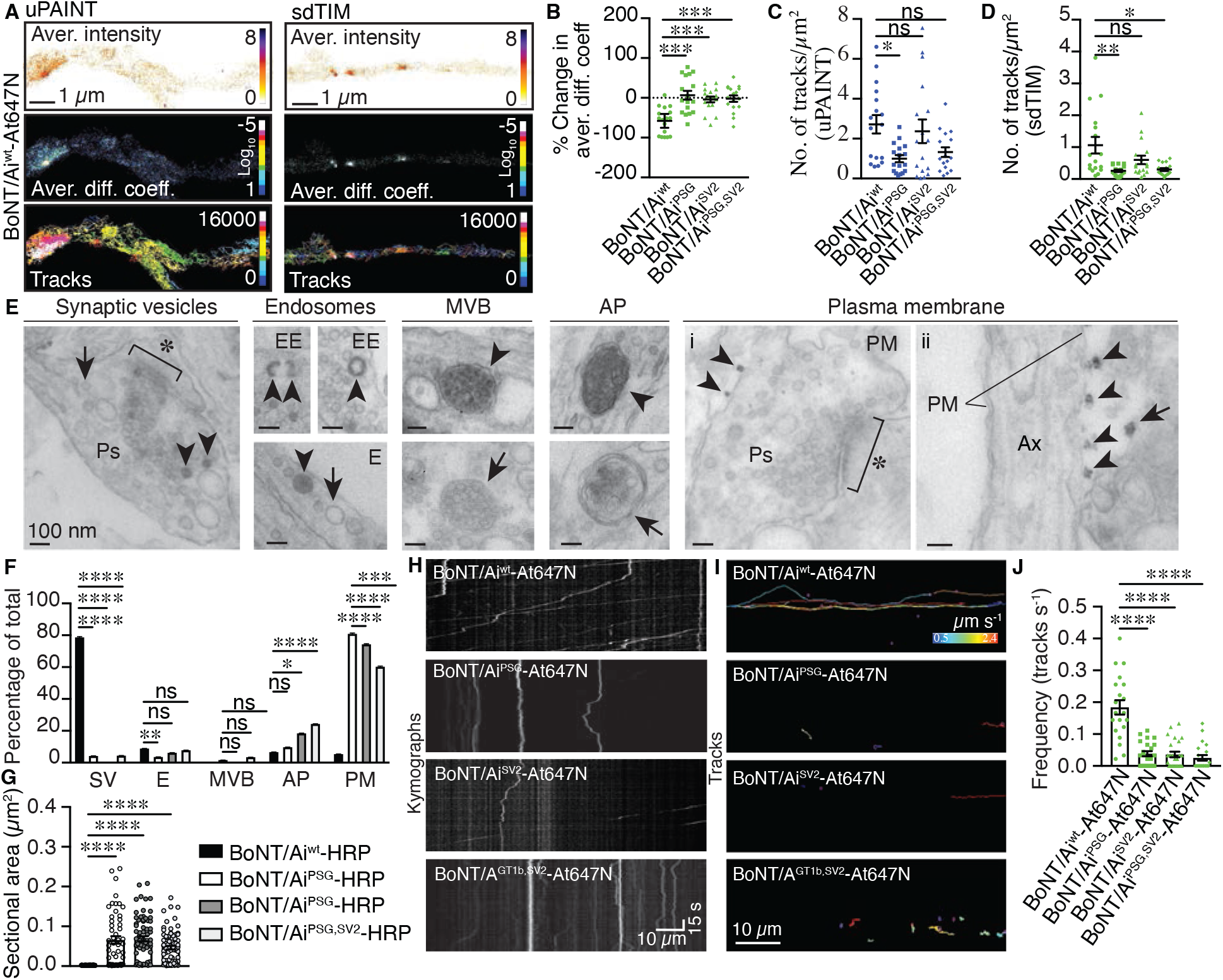
Binding to PSG and SV2 alone is insufficient for endocytic targeting of BoNT/A into synaptic vesicles. (**A**) Super-resolved average intensity (bar: high 8 to low 0 density), diffusion coefficient (bar: Log_10_D high 1 to low −5 mobility) and trajectory maps of BoNT/Ai^wt^-At647N on the plasma membrane (uPAINT) and following endocytosis (sdTIM). (**B**) Percentage change in BoNT/Ai^wt^-At647N and indicated mutant toxin mobility upon endocytosis (sdTIM) normalized to the respective mean of uPAINT. Number of BoNT/Ai^wt^-At647N and mutant trajectories (**C**) on the plasma membrane and (**D**) upon endocytosis. (**E**) EM analysis of endocytosed HRP-tagged BoNT/Ai. Representative images show cytochemically stained HRP precipitate (arrowheads). Early endosomes (EE), multivesicular bodies (MVB), structures with morphology of autophagosomes (AP), presynaptic (Ps, i) and axonal (Ax, ii) plasma membrane (PM). Postsynaptic density (asterisks), non-stained endocytic compartments or background binding on the PM (arrows) (ii) are indicated. Scale bars 100 nm. (**F**) Percentage of total BoNT/Ai^wt^-HRP and indicated mutants localized on the PM or in the indicated endocytic structures, and (**G**) the average sectional area of the HRP-stained endocytic compartments. (**H**) Kymographs and (**I**) tracks of retrograde carriers positive for the indicated holotoxins. (**J**) Frequency of retrograde carriers containing indicated toxins. (B-D) N=17 acquisitions/condition from 5-6 independent experiments, (E,F) n=60 electron micrographs/condition from 3 independent experiments, (G-I) n=28-31 axonal channels/condition from 4 independent experiments. (B-I) Non-parametric Kruskal-Wallis multiple comparison test. *p<0.05, **p<0.01, ***p<0.001, ****p<0.0001, and n.s, non-significant.

Together our results point to a model whereby BoNT/A hijacks an intrinsic Syt1-SV2 endocytic sorting mechanism by binding to a preassembled PSG-Syt1 complex and SV2, thereby inducing the formation of a tripartite PSG-Syt1-SV2 nanocluster on the neuronal plasma membrane. To visualize the tripartite PSG-Syt1-SV2 complex together with BoNT/A on the plasma membrane, we created a predictive model of the GT1b ganglioside, Syt1, and SV2 complex together with bound BoNT/A (Fig. 4 and Movie 6). The prediction was derived from the crystal structures of separate complexes, using the recently published machine learning algorithm for protein structure prediction AlphaFold2 (*28, 29*). Both SV2 and Syt1 contain significant regions of predicted structural disorder, providing flexible linkers that allow for conformational flexibility in their intracellular binding domains. Although the model is predictive, it is in line with the experimentally observed structures of GT1b, Syt1 and the luminal domain of SV2C and the tripartite PSG-Syt1-SV2 complex for selective binding and endocytic targeting of BoNT/A is structurally plausible.

**Fig. 4.**
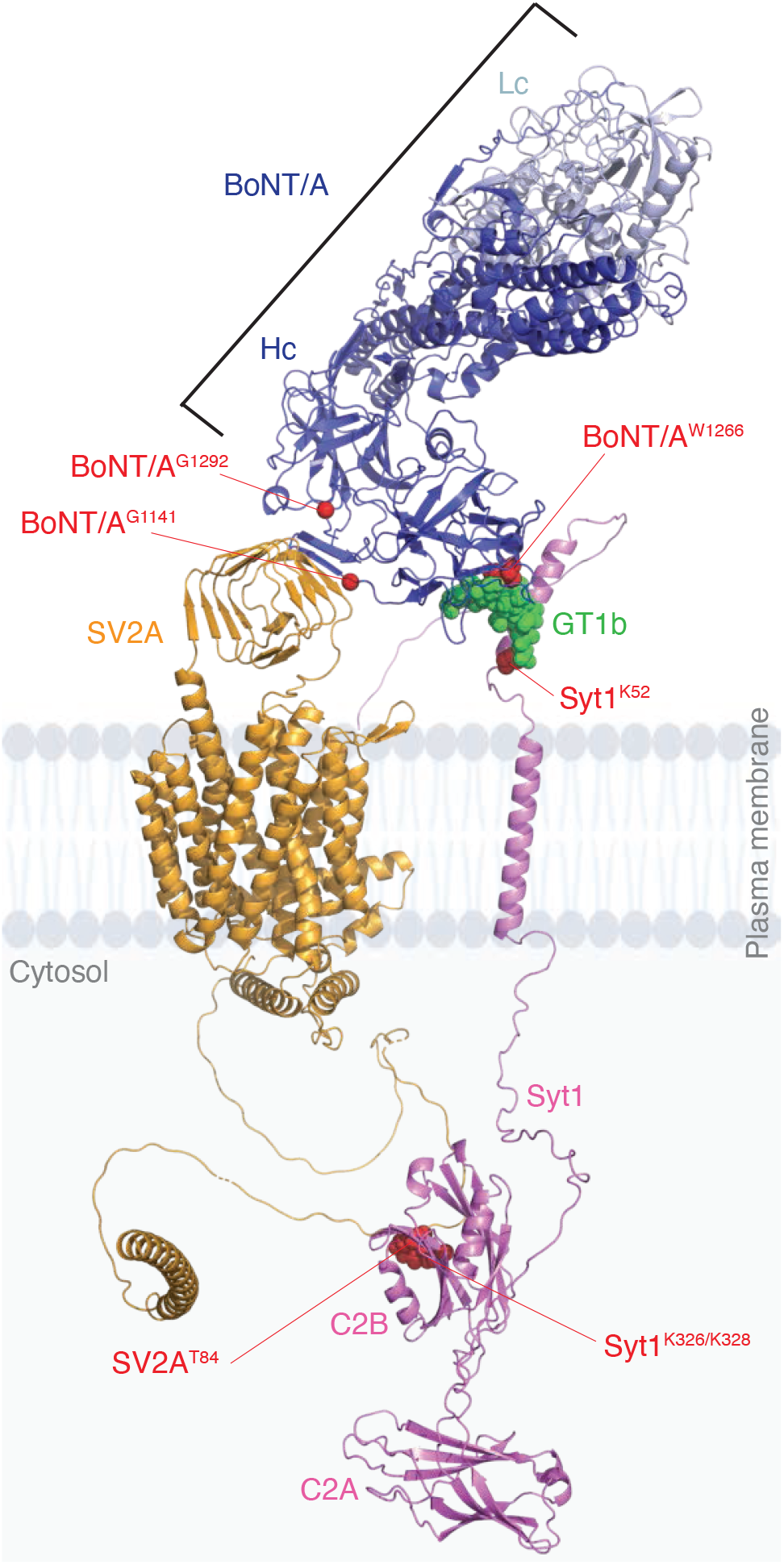
Structural model of the tripartite GT1b-Syt1-SV2 complex for selective binding and endocytic targeting of BoNT/A. Predictive model of the assembled complex between Syt1, SV2A, BoNT/A (heavy chain, H_C_, light chain, L_C_) and GT1b on the plasma membrane. Mutations used in this study are indicated as red spheres.

The connection between plasma membrane receptor nanoclustering, where specific SV proteins themselves provide a final “fail-safe” to ensure accurate retrieval of receptors and receptorbound cargo into SVs is becoming increasingly apparent (*30*). External cargoes, such as viruses and toxins, have been shown to induce nanoscale changes in the plasma membrane receptor clustering (*31*) and to target multimeric receptor clusters (*32–34*). Small changes in the number of receptors within a cluster can control whether or not a receptor and bound cargo is internalized efficiently (*35*). Our results support a model whereby BoNT/A induces Syt1-SV2 nanoclustering via coincidental binding to a PSG-Syt1 complex and SV2, subsequently triggering its own internalization into SVs to incapacitate neuronal communication. The majority of CNTs bind to two receptors, leading to irreversible plasma membrane binding (*5*). BoNT/A, B, E, F, G, and tetanus neurotoxin bind to PSGs via a conserved E(Q)…H(K/G)…SXWY…G sequence (*24*). It is therefore tempting to hypothesize a “unifying mechanism” for those CNTs that share the conserved PSG-binding site, in which the neurotoxins may also target the PSG-Syt1-SV2 complex to selectively target SVs as part of their intoxication strategy.

## Acknowledgments

We would like to thank the DNA Dream Lab (DDL) facility (HiLIFE, Institute of Biotechnology, Helsinki Finland) and personally Konstantin Kogan for the design and construction of the oligonucleotides listed in Table 1 and DNA plasmids listed in Table 2. We would like to acknowledge the facilities, and the scientific and technical assistance, of the Microscopy Australia Facility at the Centre for Microscopy and Microanalysis (CMM), especially Dr Kathryn Green, and the Queensland Brain Institute (QBI) Advanced Microscopy Facility, and the facility staff, especially Dr Rumelo Amor, at the University of Queensland. We would like to extend our thanks to Professor Claudio Rivera Baeza (Neuroscience Center, Helsinki Institute of Life Science, University of Helsinki, Finland) for his help with the interpretation of the electrophysiology experiments, Dr Ron Broide (Allergan/Abbvie) for insightful discussions and Rowan Tweedale (QBI, The University of Queensland) for critical appraisal of the manuscript.

## Author contributions

Conceptualization: MJ, AR, FAM

Formal analysis; MJ, VL, PS, JR, BMC, GB

Investigation: MJ, VL, PS, AB, JR

Methodology: MJ, VL, PS, AB, BMC, JR, RGP, SM, NK, GB, AR, FAM

Resources: MJ, VL, RG, AB, SS, MC, RC-W, JJC-W, GB, AR, FAM

Software: MJ, TW

Validation: MJ, CS, AJ

Visualization: MJ, VL, PS, JR

Supervision: MJ, RGP, AR, FAM

Writing – original draft: MJ, AR, FAM

Writing – review & editing: All authors

## Data and materials availability

Plasmids generated in this study will be deposited to Addgene. This study did not generate new unique reagents. Plasmids encoding BoNT/A and mutants are excluded for biosecurity reasons. All raw acquisitions will be deposited at The University of Queensland data repository collection and will be publicly available as of the date of publication. Any additional information required to reanalyze the data reported in this paper is available from the lead authors upon request.

