## Supplemental Material for "Synaptotagmin 1 mediates toxicity of botulinum neurotoxin type A"

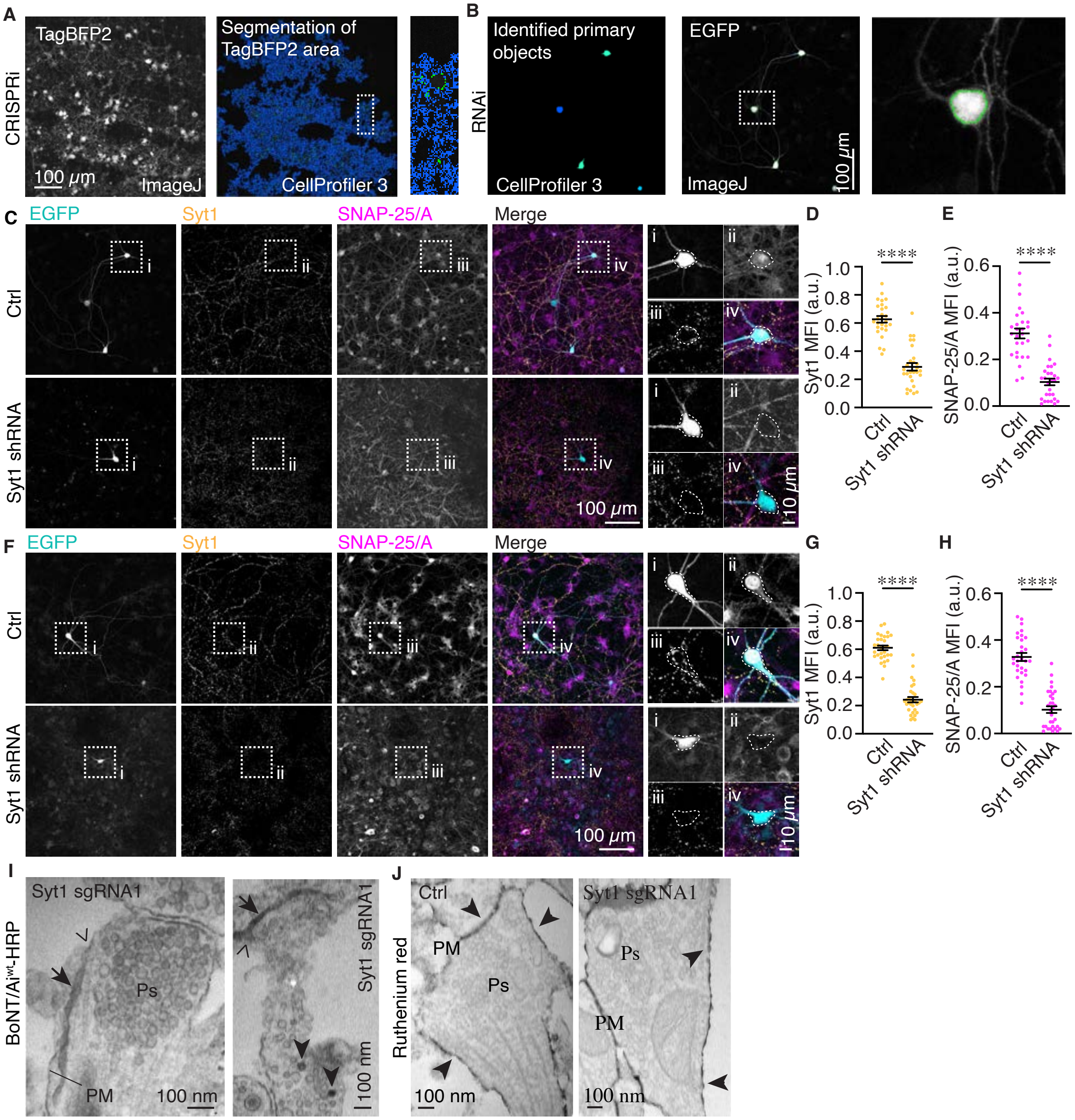

**Fig. S1: shRNA knockdown of Syt1 leads to deficient endocytic targeting of BoNT/A into synaptic vesicles and loss of BoNT/A toxic function.** (**A**) Automated segmentation analysis for quantify the mean fluorescence intensity (MFI) of Syt1 and Abobotulinumtoxin A-cleaved SNAP-25 (SNAP-25/A) levels following Syt1 KD with CRISPRi. Acquired confocal stacks were z-projected using the sum of fluorescence in ImageJ (a representative image of TagBFP2 is shown in gray on left). The 2D z-projections were then used in CellProfiler 3 to determine the area positive for TagBFP2, and to identify neurons that had received the dCas9-KRAB and the respective sgRNA. For segmentation of the TagBFP2-positive area (image in the middle), the brightest TagBFP2 fluorescent spots were identified using the Identify Primary Object tool of CP3 and the otzu algorithm (green dots; a detail from the indicated boxed area is magnified on right). Next, these primary spots were expanded following the TagBFP2 fluorescence signal until the whole TagBFP2 area of each image was detected (blue signal in the magnified image on right). MFI of the Syt1 and cleaved SNAP-25 channels were then determined within this area. (**B**) Automated segmentation analysis used to quantify the MFI of Syt1 and SNAP-25/A levels following Syt1 shRNA KD. The automated segmentation analysis was done as described above for the CRISPRi in (F), with the exception of using the EGFP fluorescent signal to identify transfected cells. The values of threshold fluorescence intensity were chosen in CellProfiler 3 so that only the brightest signal of the EGFP-positive cells was automatically detected in all images. These bright areas corresponded to the neuronal soma (boxed area is shown magnification on right), and the MFIs of Syt1 and cleaved SNAP-25/A were then quantified in these selected areas. (**C-H**) Representative Z-projection sum of a confocal stack acquired from hippocampal neurons that were co-transfected with EGFP (cyan) and either pRNAi control (ctrl) or Syt1-targeting shRNA, subjected to sdTIM uptake of 10 units of Abobotulinumtoxin A for (**C**) 30 min or (**F**) 16 h, fixed and immunostained for endogenous Syt1 (yellow) and SNAP-25/A (magenta). Boxed regions (i-iv) are magnified on right. Soma outlined with dashed line for clarity. Scatter blots of endogenous (**D**,**G**) Syt1 and (**E**,**H**) cleaved SNAP-25/A MFI in control and Syt1 shRNA-transfected hippocampal neurons following 30 min and 16 h Abobotulinumtoxin A treatment, respectively. (**I**) Representative EM images of tubular structures containing

BoNT/Ai$^{wt}$-HRP signal (arrows) in Syt1 sgRNA1 KD-transduced neurons that were observed remaining open to the extracellular space (open arrowhead). BoNT/Ai$^{wt}$-HRP in SVs (arrowheads) is indicated for reference. (**J**) EM images from control (non-targeting sgRNA) and CRISPRi Syt1 sgRNA1 transduced neurons, and stained with ruthenium red (RuR; dark electron-dense staining, arrowheads). Results are shown as average ± SEM, scatter plots indicate averages from individual acquisitions. (D,E,G,H) n=5-12/condition from 2 independent experiments. (D) Parametric unpaired *t* test, and (E,G,H) non-parametric Mann-Whitney U test. ****$p<0.0001$.

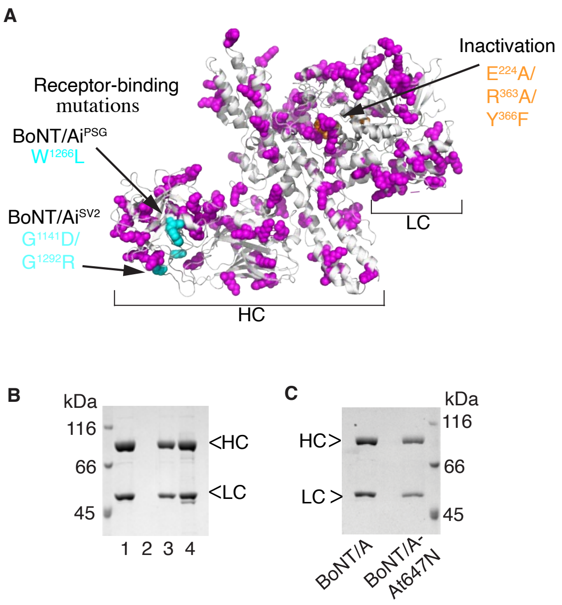

D

| Sample | Bath conc. [pM] | $t_{1/2}$ [min] | Mean $t_{1/2}$ [min] | SD [min] | CV | n |
|---|---|---|---|---|---|---|
| BoNT/A | 6 | 66 | 64.8 | 4.11 | 6.35% | 4 |
| | | 61 | | | | |
| | | 62 | | | | |
| | | 70 | | | | |
| BoNT/A-At647N | 6 | 95 | 89.7 | 8.29 | 9.24% | 6 |
| | | 81 | | | | |
| | | 88 | | | | |
| | | 82 | | | | |
| | | 103 | | | | |
| | | 89 | | | | |

E

Absorbance

| No | $A_{280}$ | $A_{647}$ | Aver. DoL |
|---|---|---|---|
| 1 | 0.692 | 0.791 | 1.46 |
| 2 | 0.719 | 0.888 | |
| 3 | 0.825 | 0.764 | |

**Fig. S2: BoNT/A-At647N toxicity in mouse phrenic nerve hemidiaphragm assay.** (**A**) Ribbon structure of BoNT/A holotoxins with lysine residues (magenta), light chain (LC) mutations (orange), and heavy chain (HC) mutations (cyan) indicated. (**B**) SDS-PAGE (10%) analysis of BoNT/A produced recombinantly in *E. coli*, yielding a solution of 2.5 µM (0.37 mg mL$^{-1}$) in PBS buffer (4.5 µL loaded in lane 1). To further concentrate the toxin, 10 nmol of BoNT/A was ultrafiltrated (MWCO 30 kDa), yielding a solution of 11.5 µM (1.7 mg mL$^{-1}$; lane 2: flow through of the ultrafiltration, and 1 µL of 1.4 ml BoNT/A solution after ultrafiltration was loaded in lane 3). Lane 4 shows 7.5 µL of the protein pellet after ultrafiltration dissolved in 140 µL (1/10) in PBS. Heavy chain, HC, light chain LC. (**C**) SDS-PAGE (10%) of BoNT/A and BoNT/A-At647N (1 µL loaded). (**D**) Functional analysis of BoNT/A and BoNT/A-At647N in the mouse hemidiaphragm assay to test the impact of the At647N labeling on BoNT/A toxicity. Employing the dose-response curve previously established for recombinant BoNT/A (*11*), the potency of BoNT/A-At647N was reduced by ~20% compared to non-labeled BoNT/A. (**E**)

Quantification of degree of labeling (DoL) of BoNT/A-At647N using absorption spectroscopy. Repeated measurements at $A_{280}$ and $A_{647}$ are shown in the table, together with the calculated average DoL. Results are shown as average ± SD, n=4-6 independent experiments (D), n=3 technical replicas (E) from one experiment.

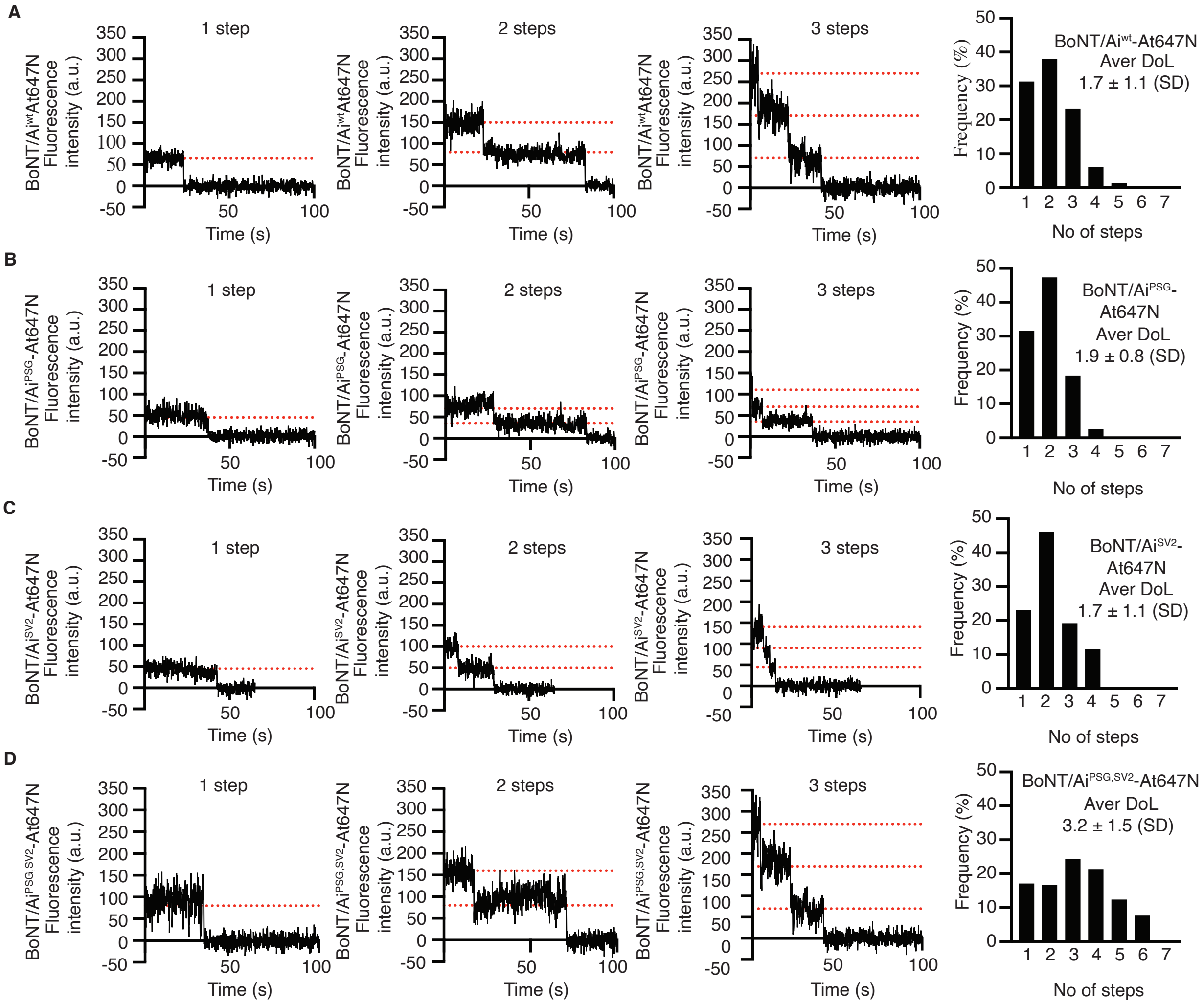

E

Degree of labelling

| No | Sample | $A_{280}$ | $A_{647}$ | Aver DoL |
|---|---|---|---|---|
| 1 | BoNT/Ai$^{wt}$-At647N | 0.400 | 0.465 | 1.68 |
| 2 | BoNT/Ai$^{wt}$-At647N | 0.360 | 0.454 | |
| 3 | BoNT/Ai$^{wt}$-At647N | 0.320 | 0.415 | |
| 4 | BoNT/Ai$^{wt}$-At647N | 0.300 | 0.415 | |

| No | Sample | $A_{280}$ | $A_{647}$ | Aver DoL |
|---|---|---|---|---|
| 1 | BoNT/Ai$^{PSG}$-At647N | 0.280 | 0.446 | 1.83 |
| 2 | BoNT/Ai$^{PSG}$-At647N | 0.330 | 0.420 | |
| 3 | BoNT/Ai$^{PSG}$-At647N | 0.280 | 0.397 | |

| No | Sample | $A_{280}$ | $A_{647}$ | Aver DoL |
|---|---|---|---|---|
| 1 | BoNT/Ai$^{SV2}$-At647N | 0.240 | 0.388 | 2.17 |
| 2 | BoNT/Ai$^{SV2}$-At647N | 0.260 | 0.374 | |
| 3 | BoNT/Ai$^{SV2}$-At647N | 0.220 | 0.396 | |

| No | Sample | $A_{280}$ | $A_{647}$ | Aver DoL |
|---|---|---|---|---|
| 1 | BoNT/Ai$^{PSG,SV2}$-At647N | 0.450 | 1.403 | 3.50 |
| 2 | BoNT/Ai$^{PSG,SV2}$-At647N | 0.460 | 1.085 | |
| 3 | BoNT/Ai$^{PSG,SV2}$-At647N | 0.480 | 1.088 | |

**Fig. S3: Quantification of the degree of At647N-labeling of BoNT/Ai holotoxins.**

Representative single-molecule fluorescence emission traces of (**A**) BoNT/Ai$^{wt}$-At647N, (**B**)

BoNT/Ai$^{PSG}$-At647N, (**C**) BoNT/Ai$^{SV2}$-At647N, and (**D**) BoNT/Ai$^{PSG,SV2}$-At647N recorded on a custom-made flow-chamber glass-bottom. The frequency (%) distribution of emission step numbers is shown in the bar graphs on right with the respective average DoL for each toxin. (**E**) Absorption spectroscopy recordings of indicated toxins. Repeated measurements of $A_{280}$ and $A_{647}$ are shown in the table together with calculated average DoL for each toxin. Results are shown as average ± SD, n=163 BoNT/Ai$^{wt}$-At647N (A), n=38 BoNT/Ai$^{PSG}$-At647N (B), n=78 BoNT/Ai$^{SV2}$-At647N (C), n=234 BoNT/Ai$^{SV2}$-At647N (D) and n=3 technical replicas (E) all from one experiment/condition.

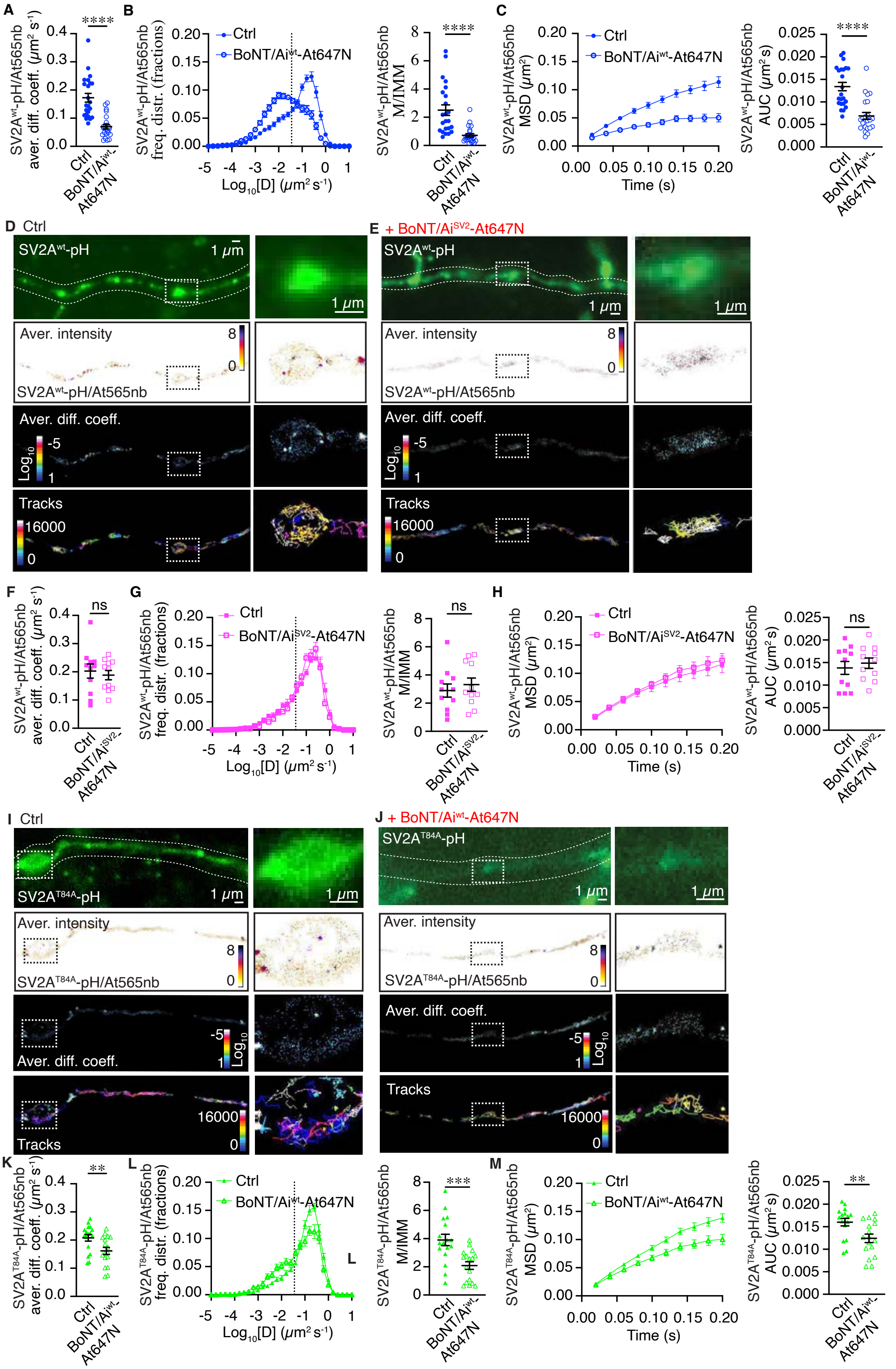

A
SV2A^wt-pH/At565nb aver. diff. coeff. (μm² s⁻¹)
****
Ctrl
BoNT/Ai^wt-At647N
B
SV2A^wt-pH/At565nb freq. distr. (fractions)
Ctrl
BoNT/Ai^wt-At647N
Log10[D] (μm² s⁻¹)
SV2A^wt-pH/At565nb M/IMM
C
SV2A^wt-pH/At565nb MSD (μm²)
Time (s)
SV2A^wt-pH/At565nb AUC (μm² s)
D Ctrl
SV2A^wt-pH
1 μm
Aver. intensity
SV2A^wt-pH/At565nb
Aver. diff. coeff.
Log10
Tracks
16000
E + BoNT/Ai^SV2-At647N
F
ns
Ctrl
BoNT/Ai^SV2-At647N
G
H
I Ctrl
SV2A^T84A-pH
SV2A^T84A-pH/At565nb
J + BoNT/Ai^wt-At647N
K
SV2A^T84A-pH/At565nb aver. diff. coeff. (μm² s⁻¹)
**
L
SV2A^T84A-pH/At565nb freq. distr. (fractions)
SV2A^T84A-pH/At565nb M/IMM
***
M
SV2A^T84A-pH/At565nb MSD (μm²)
SV2A^T84A-pH/At565nb AUC (μm² s)

**Fig. S4: BoNT/Ai-induced changes to the SV2 mobility on the plasma membrane depend on Syt1-SV2 interactions.** Single-molecule uPAINT mobility of SV2A$^{wt}$-pH/At565nb as (**A**) average diffusion coefficient, (**B**) frequency distribution of diffusion coefficient and mobile-to-immobile (M/IMM) ratio, (**C**) mean square displacement (MSD) and area under the MSD curve (AUC) in the absence (ctrl; high $K^+$) and presence of BoNT/Ai$^{wt}$-At647N (toxin diluted in high $K^+$). (**D**,**E**) Top panel shows low resolution images of neurons expressing SV2A$^{wt}$-pH. uPAINT imaging was performed in (**D**) control conditions (high $K^+$) and (**E**) following BoNT/Ai$^{SV2}$-At647N treatment (toxin diluted in high $K^+$) and the resulting average intensity, diffusion coefficient and trajectory maps of SV2A$^{wt}$-pH/At565nb are indicated. Boxed areas magnified on right. (**F-H**) Single-molecule uPAINT mobility of SV2A$^{wt}$-pH/At565nb in the absence (ctrl; high $K^+$) and presence of BoNT/Ai$^{SV2}$-At647N (toxin diluted in high $K^+$). (**I**,**J**) Top panel shows low resolution images of neurons expressing SV2A$^{T84A}$-pH. uPAINT imaging was performed in (**I**) control conditions (high $K^+$) and (**J**) following BoNT/Ai$^{SV2}$-At647N treatment (toxin diluted in high $K^+$) and the resulting average intensity, diffusion coefficient and trajectory maps of SV2A$^{T84A}$-pH/At565nb are indicated. (**K-M**) Single-molecule uPAINT mobility of SV2A$^{T84A}$-pH/At565nb in the absence (ctrl; high $K^+$) and presence of BoNT/Ai$^{wt}$-At647N (toxin diluted in high $K^+$). Results are shown as average ± SEM, scatter plots indicate averages from individual acquisitions. N=11-23 acquisitions/condition from 3-5 independent experiments. Non-parametric Mann-Whitney U test (A-C,H), parametric unpaired t test (F,G,L,M). $^{**}p<0.01$, $^{***}p<0.001$, $^{****}p<0.0001$, and n.s, non-significant.

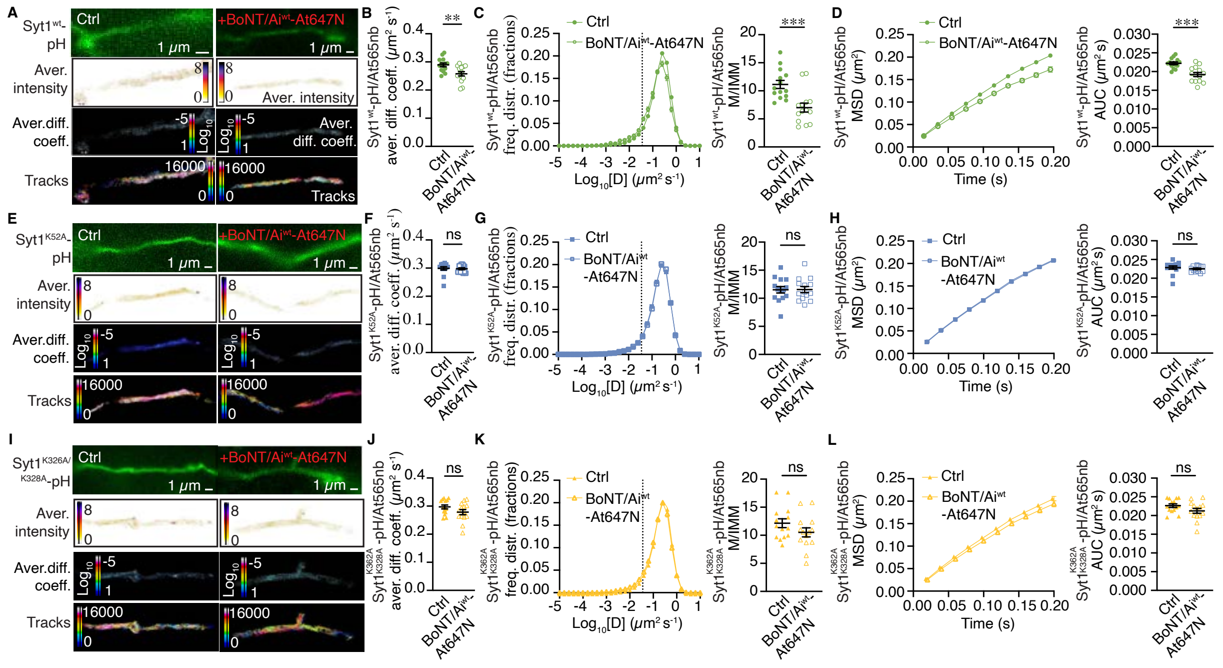

**Fig. S5: BoNT/Ai-induced changes on Syt1 mobility on the plasma membrane depend on Syt1-SV2 interactions.** (**A**) Top panel shows low resolution images of neurons expressing Syt1$^{wt}$-pH. uPAINT imaging was performed in control conditions (high K$^{+}$) and following BoNT/Ai$^{wt}$-At647N treatment (toxin diluted in high K$^{+}$) and the resulting average intensity, diffusion coefficient and trajectory maps of Syt1$^{wt}$-pH/At565nb are indicated. Single-molecule uPAINT mobility of Syt1$^{wt}$-pH/At565nb as (**B**) average diffusion coefficient, (**C**) frequency distribution of diffusion coefficient and mobile-to-immobile (M/IMM) ratio, (**D**) mean square displacement (MSD) and area under the MSD curve (AUC) in the absence (ctrl; high K$^{+}$) and presence of BoNT/Ai$^{wt}$-At647N (toxin diluted in high K$^{+}$). (**E**) Top panel shows low resolution images of neurons expressing Syt1$^{K52A}$-pH. uPAINT imaging was performed in control conditions (high K$^{+}$) and following BoNT/Ai$^{wt}$-At647N treatment (toxin diluted in high K$^{+}$) and the resulting average intensity, diffusion coefficient and trajectory maps of Syt1$^{K52A}$-pH/At565nb are indicated. Boxed areas magnified on right. (**F-H**) Single-molecule uPAINT mobility of Syt1$^{K52A}$-pH/At565nb in the absence (ctrl; high K$^{+}$) and presence of BoNT/Ai$^{wt}$-At647N (toxin diluted in high $K^+$). (**I**,**J**) Top panel shows low resolution images of neurons expressing $Syt1^{K326A,K328A}$-pH. uPAINT imaging was performed in control conditions (high $K^+$) and following BoNT/$Ai^{wt}$-At647N treatment (toxin diluted in high $K^+$) and the resulting average intensity, diffusion coefficient and trajectory maps of $Syt1^{K326A,K328A}$-pH/At565nb are indicated. (**K-M**) Single-molecule uPAINT mobility of $Syt1^{K326A,K328A}$-pH/At565nb in the absence (ctrl; high $K^+$) and presence of BoNT/$Ai^{wt}$-At647N (toxin diluted in high $K^+$). Results are shown as average ± SEM, scatter plots indicate averages from individual acquisitions. n=11-19/condition from 3-5 independent experiments. Non-parametric Mann-Whitney U test (F,H,L), parametric unpaired *t* test (B-D,G,K,J). **p<0.01, ***p<0.001, and n.s, non-significant.

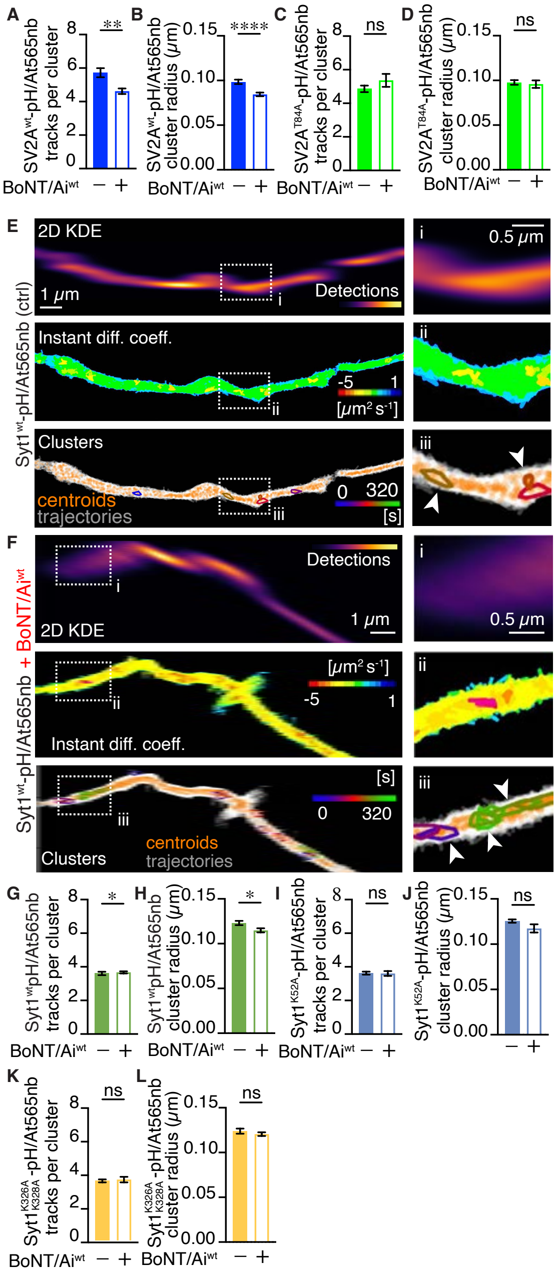

**Fig. S6: BoNT/Ai-induced changes in SV2 and Syt1 receptor clustering.** NASTIC quantification of (**A**,**B**) SV2A$^{wt}$-pH/At565nb and (**C**,**D**) SV2A$^{T84A}$-pH/At565nb nanoclustering in control conditions (high K$^{+}$;"-") and following BoNT/Ai$^{wt}$-At647N treatment (toxin diluted in high K$^{+}$ "+"). Nanocluster analysis of Syt1$^{wt}$-pH/At565nb in (**E**) control condition (high K$^{+}$) and (**F**) following high K$^{+}$ stimulation with BoNT/Ai$^{wt}$-At647N. Boxed areas (i-iii) magnified on right, arrowheads indicate nanoclusters. NASTIC quantification of (**G**,**H**) Syt1$^{wt}$-pH/At565nb,

(**I**,**J**) Syt1$^{K52A}$-pH/At565nb and (**K**,**L**) Syt1$^{K326AA,K328A}$-pH/At565nb nanoclustering in control conditions (high $K^+$;"-") and following BoNT/Ai$^{wt}$-At647N treatment (toxin diluted in high $K^+$ "+"). Results are shown as average±SEM, n=11-19 acquisitions/condition from 3-5 independent experiments. Parametric unpaired t test (B, D, J, L), and non-parametric Mann-Whitney U test (A, C, G, H, I, K). *p<0.05, **p<0.01, ****p<0.0001, and n.s, non-significant.

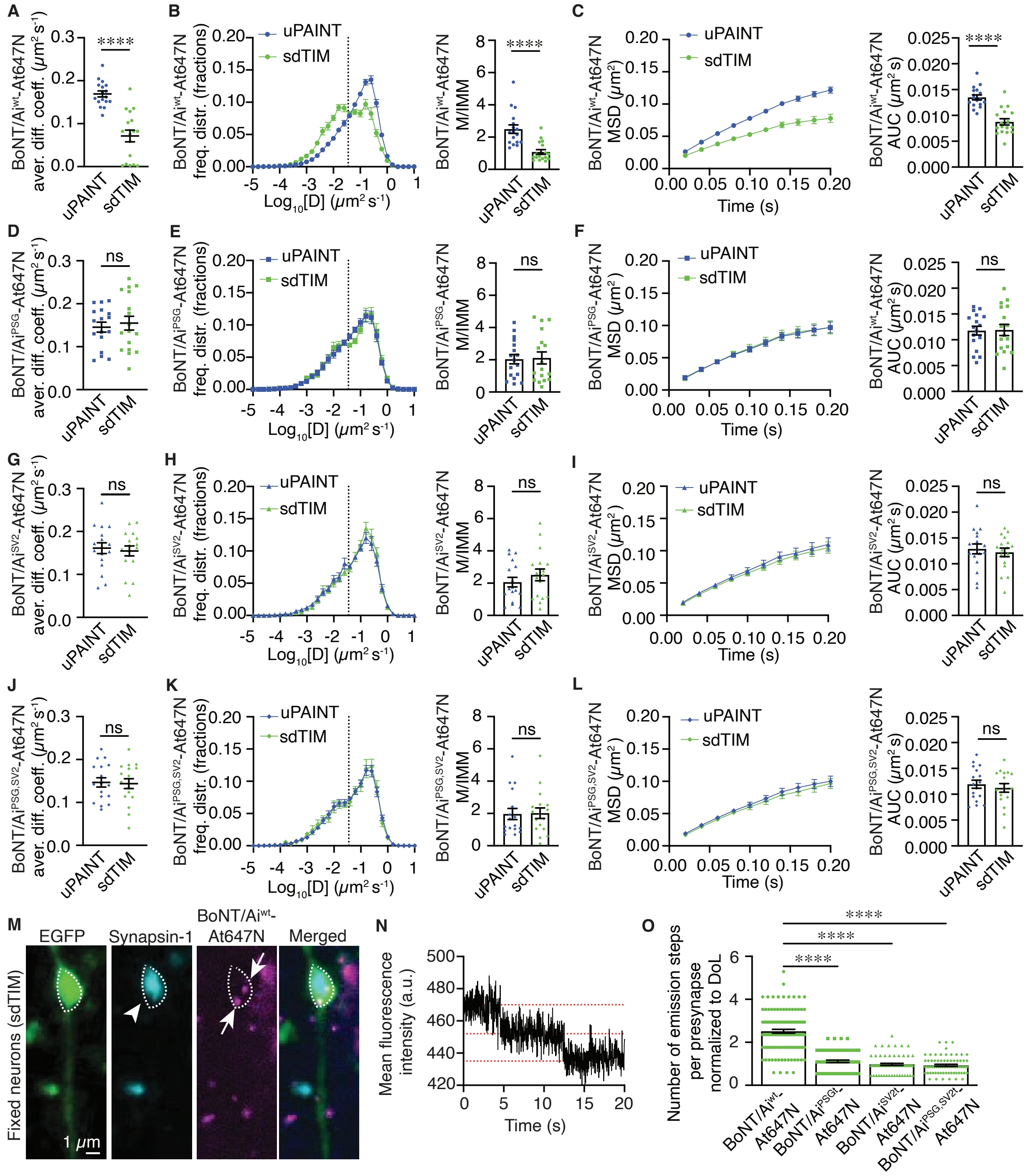

**Fig. S7: Dynamic nanoscale organization of BoNT/Ai holotoxin upon binding and internalization in neurons.** Single-molecule uPAINT and sdTIM mobility quantification of (**A**-**C**) BoNT/Ai$^{wt}$-At647N, (**D**-**F**) BoNT/Ai$^{PSG}$-At647N, (**G**-**I**) BoNT/Ai$^{SV2}$-At647N and (**J**-**L**) BoNT/Ai$^{PSG,SV2}$-At647N indicated as average diffusion coefficient, frequency distribution (fractions) of $Log_{10}$ diffusion coefficient, mobility to immobile fraction (M/IMM), mean square displacement (MSD) and area under the MSD curve (AUC). (**M**) Representative image of an axonal segment from a hippocampal neuron expressing EGFP (green), subjected to sdTIM uptake of BoNT/$Ai^{wt}$-At647N (magenta, arrows), followed by fixation and immunostaining of endogenous synapsin-1 (cyan, arrowhead). (**N**) Fluorescence emission of BoNT/$Ai^{wt}$-At647N from the fixed neurons was recorded and the representative trace shows the fluorescence emission (2 steps) of internalized BoNT/$Ai^{wt}$-At647N in a presynaptic ROI (synapsin-1 signal; outlined with dashed line in J). (**O**) Scatter plots of the number of emission steps of internalized BoNT/$Ai^{wt}$-At647N and mutants in presynaptic ROIs, normalized to the degree of labeling (DoL) of the respective holotoxin. Results are shown as average ± SEM, dots in scatter plots indicate averages from individual acquisitions (A-L) or presynapses (O). N=17/condition from 5-6 independent experiments (A-L), n=71-116 presynapses/condition from 3 independent experiments (M-O). Non-parametric Mann-Whitney U test (B,E,K), parametric unpaired *t* test (A,C,D,F,G,H-J,L), non-parametric Kruskal-Wallis multiple comparison test (O). ****$p<0.0001$, and n.s, non-significant.

### Materials and Methods

#### Materials

#### Hippocampal neurons

Experiments were conducted with the approval of The University of Queensland Animal Ethics Committee (Approval #2016/AE000254 and 2019/AE000244). Pregnant Sprague Dawley rats (Charles River Laboratories; https://www.criver.com) were individually housed and maintained in a dedicated animal husbandry area at room temperature ~22ºC in pathogen-controlled laboratory conditions. Rats were kept on a 12:12 light/dark cycle and received water and food *ad libitum*. All experiments were conducted on hippocampal neurons obtained from rats at embryonic day 18 (E18) as previously described (*1*). Both male and female embryos were used for all experiments. Dissected hippocampal neurons were cultured in neuronal culture medium, composed of Neurobasal medium (Gibco-Thermo Fisher, Cat. #12348-017), 1x serum-free B27 supplement (Gibco-Thermo Fisher, Cat. #17504-044), 1x GlutaMAX supplement (Gibco-Thermo Fisher, Cat. #35050-061) and a penicillin (100 U $mL^{-1}$)/streptomycin (100 µg $mL^{-1}$, Invitrogen-Thermo Fisher, Cat. #15140-122) mix as previously described (*1*), on poly-L-lysine–coated (Sigma-Aldrich, Cat. #P2636) glass-bottom dishes at 80,000 (CellVis, Cat. # D29-20-1.5-N for all super-resolution imaging), and 30,000-50,000 (CellVis, Cat. #D29-10-1.5-N for proximity ligation assay, CRISPR interference and electron microscopy) cells per coverglass. For electrophysiology, hippocampal neurons were plated in 50,000 cells per poly-L-lysine–coated 5-mm coverslip placed in D29-10-1.5-N glass-bottom wells. All transfections were performed on day *in vitro* 14 (DIV14), and on DIV7 for shRNA Syt1 knockdown (KD) experiments, using Lipofectamine 2000 reagent (Invitrogen, Cat. #11668019) according to the manufacturer's instructions. CRISPRi lentiviral transduction was performed on DIV14 (for 7 days) and subsequent rescue experiments on DIV19 (for 48 h). All live-cell imaging and sample processing were performed on mature neurons on DIV21-22.

**Antibodies and reagents**

Anti-SV2A antibody (Cat. #ab32942) and anti-synaptotagmin-1 antibody (Cat. #ab13259) were purchased from Abcam. Anti-SV2C antibody (Cat. #119203), anti-synapsin-1 (Cat. # 06 011) and anti-GFP sdAb antibody - FluoTag-Q At565 (Cat. #N0301-At565-S) were all from Synaptic Systems. Anti-SNAP-25/A was a kind gift from D. Sesardic Division of Bacteriology, National Institute for Biological Standards and Control, United Kingdom. ATTO 647N NHS-ester (Cat. #AD 647N-31) was from ATTO-TEC GmbH.

**Mammalian expression constructs**

The pHluorin-Syt1 K52A (Syt1$^{K52A}$-pH) construct was created by site-directed mutagenesis as per manufacturer's instructions (QuikChange Lightning Site-Directed Mutagenesis Kit, Agilent Technologies, Cat. #210518). The mutagenesis primers used were Forward: 5' - GTTTATGAATGAGCTGCATGCAATTCCATTGCCACCGTG 3' and Reverse: 5'-CACGGTGGCAATGGAATTGCATGCAGCTCATTCATAAAC 3'. The construct was sequenced in the Australian Genome Research Facility at The University of Queensland. Synaptotagmin1 shRNA was inserted into pSUPER-neo+GFP (with GFP replaced by mCerulean(*2*)). pSuper-neo+mCerulean was cut with BgIII and XhoI to insert annealed shRNA oligonucleotides by T4 ligation; the BgIII site is destroyed by this process. Sense oligonucleotide 5' - GATCCCCGAGCAAATCCAGAAAGTGCAATTCAAGAGATTGCACTTTCTGGATTTGCT

CTTTTTC 3’ and antisense oligonucleotide 5’ TCGAGAAAAA GAGCAAATCCAGAAAGTGCAA TCTCTTGAATTGCACTTTCTGGATTTGCTC GGG 3’.

**Lentiviral CRISPRi and rescue plasmids**

In the CRISPRi system, the nuclease null form Cas9 (dCas9) was fused to three different single guide RNAs (sgRNA1-3) and to the transcription repressor KRAB (dCas9-KRAB) to block Syt1 gene transcription. This technique avoids the use of insertions or deletions (indels) which guarantees the genotypic consistency of neurons following gene suppression, thereby making CRISPRi particularly suitable for Syt1 KD in the neurons, as demonstrated previously (*3*). All sgRNAs for rat Syt1 were constructed on the same backbone vector (#AAAA-0244, termed ‘dCas9-KRAB-TagBFP2’; Table S2) by DNA Dream Lab. We used pLV hU6-sgRNA hUbC-dCas9-KRAB-T2a-Puro construct (dCAS9-KRAB; a gift from Charles Gersbach, Addgene plasmid #71236;http://n2t.net/addgene:71236; RRID: Addgene_71236 (*4*)), where the puromycin-resistance gene was replaced by the TagBFP2 (blue fluorescent protein 2) marker, which was PCR-amplified from the ACE2 g1 + Cas9 plasmid (a gift from Jason Sheltzer, Addgene plasmid #153011; http://n2t.net/addgene:153011; RRID:Addgene_153011). The replacement was made with the NEBuilder kit (NEB, Cat. #E2621S) by concatenating the PCR-linearized dCAS9-KRAB construct and TagBFP2 fluorescent marker with the primers indicated in Table S1. The final backbone expresses gRNA under the human U6 promoter, with dCAS9-KRAB being fused with self-cleaving peptide (via T2A) and TagBFP2 fluorescent marker, - all under the UbC promoter. To find the targets for gRNAs on the 5’-UTR of the rat Syt1 gene, we used Chop-Chop (*5-7*) (https://chopchop.cbu.uib.no) and CRISPick (*8, 9*) (aka GPP; https://portals.broadinstitute.org/gppx/crispick/public) and Benchling softwares (Benchling, https://www.benchling.com/molecular-biology/). We selected 3 locations manually and designed oligonucleotides with CACC and AAAC overhangs on the 5' position of the forward and reverse primers, respectively, for cloning into the dCas9-KRAB-TagBFP2 vector. Each pair of primers was annealed and ligated into the BsmBI-linearized dCas9-KRAB-TagBFP2 vector following the manufacturer's instructions. The resulting plasmids are listed in Table S2 (Syt1 sgRNA1 #AAAA-0245, Syt1 sgRNA1 #AAAA-0246, and Syt1 sgRNA1 #AAAA-0247). All the sequences were confirmed by Sanger sequencing to match the original design. For Syt1 rescue experiments, the pLenti6.3-Syt1$^{wt}$-pH vector (#AAAA-0238; Table S2) with N-terminal pHluorin-tag was produced by assembling the PCR-linearized *Rattus norvegicus* Syt1 gene under the human synapsin1 promoter as one DNA fragment from the pSUPER-pHluorin-Syt1 (Syt1$^{wt}$-pH) plasmid and PCR-linearized pLenti6.3 vector, using NEBuilder HiFi DNA Assembly Master Mix (NEB, Cat. #E2621), following the standard manufacturer's protocol. The existing CMV promoter in the original pLenti6.3 was eliminated during the PCR step. We used KAPA HiFi HotStart ReadyMix (Roche, Cat. #7958927001) DNA polymerase for the PCR reactions according to the manufacturer's protocol.

**Production of inactivated BoNT/Ai holotoxins**

Enzymatically inactivated, full-length BoNT/Ai$^{wt}$ (E224A/R363A/Y366F) (*10*) and the respective receptor-binding mutant BoNT/Ai$^{PSG}$ (E224A/R363A/Y366F + W1266L), BoNT/Ai$^{SV2}$ (E224A/R363A/Y366F + G1141D/G1292R) and BoNT/Ai$^{PSG\text{-}SV2}$ (E224A/R363A/Y366F + W1266L/G1141D/G1292) holotoxin were produced recombinantly in the *E. coli* BL21 DE3 strain (NEB, Cat. #C2527H) in the laboratory of Dr. Andreas Rummel. All mutations were generated by two-step PCR and verified by DNA sequencing. Inactive BoNT/Ai$^{wt}$ and mutants carrying a C-terminal His6-tag were purified on $Co^{2+}$-Talon matrix (Takara Bio Europe S.A.S.) and eluted with 50 mM Tris-HCl (pH 8.0), 150 mM NaCl, and 250 mM imidazole. For proteolytic activation and removal of the affinity tag, BoNTs were incubated for 16 h at room temperature with 0.01 U bovine thrombin (Sigma-Aldrich Chemie GmbH) per µg of BoNT. Subsequent gel filtration (Superdex-200, Cytiva) was performed in phosphate buffered saline (PBS, pH 7.4). For storage, enzymatically inactivated, full-length BoNT/$A^{wt}$ and mutants were snap-frozen in liquid nitrogen and maintained at -80°C.

**Production of active BoNT/A for mouse phrenic nerve hemidiaphragm assay**

Wild-type active BoNT/A was produced recombinantly in BL21(DE3) *E. coli* ((NEB, Cat. #C2527H) as previously described (*11*) under biosafety level 2 containment (project number GAA A/Z 40654/3/123), yielding a solution of 2.5 µM (0.37 mg $mL^{-1}$) BoNT/A in PBS buffer. To obtain a high degree of labeling (DoL; manufacturer recommended protein concentrations > 1 mg $mL^{-1}$), 10 nmol of BoNT/A were concentrated by ultrafiltration (MWCO 30 kDa) by a factor of ~4 yielding a solution of 11.5 µM (1.7 mg $mL^{-1}$) BoNT/A.

**At647N labeling of active and inactivated BoNT/A and determination of the degree of labeling**

To map the potential At647N-labeling sites in the toxin structure, the BoNT/A lysine (L) residues, together with point mutations inactivating the catalytic activity (E224A/R363A/Y366F) and receptor binding mutations (W1266L/G1141D/G1292R), were indicated in the ribbon structure of BoNT/A holotoxin using the Pymol program (Schrodinger, https://pumol.org/2/)(fig. S2A). This demonstrated a homogenous distribution of the 103 lysine residues throughout the BoNT/A structure, and we assume a random coupling of At647N to these residues. BoNT/Ai$^{wt}$, BoNT/Ai$^{W1266L}$ (BoNT/Ai$^{PSG}$), BoNT/Ai$^{G1141D/G1292}$ (BoNT/Ai$^{SV2}$), and BoNT/Ai$^{W1266L/G1141D/G1292}$ (BoNT/Ai$^{PSG,SV2}$) holotoxins were labeled with Atto 647N NHS-ester (At647N; ATTO-TEC Gmbh, Cat. #AD 647N-31) according to the manufacturer's protocol. Briefly, At647N was dissolved in water-free DMSO and the pH of BoNT/Ai dilution in PBS buffer was adjusted to 8.3 with 0.2 M $NaHCO_3$. A 3:1 (molar) ratio of At647N-dye:BoNT/Ai was mixed, shielded from light and incubated for 1.5 h at room temperature in slow rotation. The non-bound dye was then removed using Sephadex G-25 columns (Cytiva, Cat. #C985C42) and the first blue fraction containing the labeled At647N-BoNT/A was collected, aliquoted and stored at -80°C.

The degree of labelling (DoL) of BoNT/Ai$^{wt}$-At647N, and the receptor-binding mutants, was first quantified in a cell-free flow-chamber assay by quantifying the number of At647N fluorescence emission steps over time, as previously described (*1, 12*). The average number of emission steps was 1.7 ± 1.1 (SD) for BoNT/Ai$^{wt}$-At467N, 1.9 ± 0.8 for BoNT/Ai$^{PSG}$-At467N, 1.7 ± 1.1 for BoNT/Ai$^{SV2}$-At467N, and 3.2 ± 1.5 for BoNT/Ai$^{PSG,SV2}$-At467N, indicating that each BoNT/A molecule was labeled with one to three At647N molecules. The DoL was then confirmed using a NanoDrop ND 1000 (Thermo Scientific) spectrophotometer (fig. S3E) using the equation (where $A_{280}$ is the absorption maximum of BoNT/A, and $A_{647}$ is the absorption maximum of At647N):

$$DoL = (A647 \times \frac{\varepsilon BoNTA}{(A280 - A647 \times CF280) \times \varepsilon max})$$

The extinction coefficient of inactivated BoNT/A at 280 nm (εBoNTA) was calculated as follows (where n(W), n(Y) and n(C) are the number of tryptophans, tyrosines and cysteines in the BoNT/A amino acid sequence, respectively):

$$\mathrm{\varepsilon BoNTA} = \mathrm{n(W)} \times 5500 + \mathrm{n(Y)} \times 1490 + \mathrm{n(C)} \times 125 \times 1/\mathrm{M} \times \mathrm{cm}$$

Using Nanodrop ND 1000's Proteins & Labels test type, a new dye At647N with the following specifications was created: $CF_{260}$(At647N)= 0.04; $CF_{280}$(At647N)= 0.03; $\lambda_{abs}$ = 647 nm; $\varepsilon_{max}$ = 1.5 * $10^5$ * 1/($M^{-1}$ $cm^{-1}$). The sample type was set to other protein (E 1%) BoNT/A $\varepsilon^{1\%}$ = ($\varepsilon_{BoNT/A}$ * 10) / $M_w$ = 192395 / 150000 = 12.83. The absorption was measured at 280 nm and 647 nm from three to four technical replicates, with the estimated average DoL being 1.68 for BoNT/$Ai^{wt}$-At647N, 1.83 for BoNT/$Ai^{PSG}$-At467N, 2.17 for BoNT/$Ai^{SV2}$-At467N, and 3.5 for BoNT/$Ai^{PSG,SV2}$-At467, supporting the results obtained from the cell-free flow-chamber assay.
At647N-labeling of active BoNT/A was performed as described above, and analysed with SDS-PAGE along with non-labeled active BoNT/A, demonstrating no apparent change in the size of the toxins following the labeling (fig. S2C). The BoNT/A-At647N DoL was then determined using spectrophotometer NanoDrop 1000 as described above (fig. S2E), indicating that on average BoNT/A-At647N contained one or two At647N-fluorophores (average DoL 1.46).

**HRP labeling of holotoxins**

A Lightning-Link HRP Antibody Labeling Kit (Novus Biologicals, Cat. #701-0000) was used to conjugate the inactivated BoNT/$Ai^{wt}$, BoNT/$Ai^{PSG}$, BoNT/$Ai^{SV2}$ and BoNT/$Ai^{PSG,SV2}$ holotoxins with HRP. For the random direct conjugation of HRP to any available amine group in the toxin structure, the labeling was performed according to the manufacturer's instructions in a 1:4 molar ratio of holotoxin:HRP. All conjugated toxins were stored at −20°C with the addition of 50% glycerol as a stabilizer.

**Lentiviral production**

For lentiviral production, 5.0x10$^{6}$ Lenti-XTM 293T Cell Line (Takara Bio, Cat. #632180) cells were seeded into T175 flasks (Nunc, Cat. #159910) in 35 mL of Dulbecco's Modified Eagle Medium (DMEM) with 4.5 g $L^{-1}$ glucose (Gibco, Cat. #11995-065), 10% fetal bovine serum (FBS; Gibco, Cat. #26140-079), 100 U $mL^{-1}$ Penicillin/streptomycin (Invitrogen-Thermo Fisher, Cat #15140-122), a 1x L-glutamine (Gibco, Cat. # 25030). The following day, 60% to 70% confluent cells were transfected with Transit LT-1 (Mirus, Cat. #2304) according to the manufacturer´s instructions. Briefly, 150 µL of Transit LT-1 was diluted into 6 mL of Opti-MEM (Gibco, Cat. #31985-070) and 50 µg (total) of the following plasmids were separately diluted in Opti-MEM: pMDLg/pRRE (a gift from Didier Trono, Addgene plasmid #12251; http://n2t.net/addgene:12251; RRID:Addgene_12251(*13*)), pRSV-REV (a gift from Didier Trono, Addgene plasmid #12253; http://n2t.net/addgene:12253; RRID:Addgene_12253(*13*)), pMD2.G (a gift from Didier Trono, Addgene plasmid #12259; http://n2t.net/addgene:12259; RRID:Addgene_12259), and either the CRISPRi plasmid encoding TagBFP2 and the dCas9-KRAB, or one of the sgRNA1-3 guides, or rescue plasmid encoding pLenti6.3-Syt1$^{wt}$-pH (see Table S2). The plasmid ratio for pMDL g/p RRE, pRSV-REV, pMD2-G and the CRISPRi/rescue constructs was 1:1:0.5:2.5, respectively. After 5 min, the two solutions where mixed and incubated for 20 min at room temperature. The mix was then added to the cells drop by drop and the cells were incubated for 8 h at 37ºC in 5% $CO_2$ atmosphere, after which the culture medium was replaced with complete growth medium (without antibiotics). 48 h after transfection, the medium was collected and centrifuged at 3,800 x *g* for 15 min at 4ºC. The supernatant was then centrifuged at 100,000 x *g* in a SW32Ti rotor for 2 h at 4ºC, over a 4.5 mL cushion of 20% sucrose prepared in Neurobasal medium. The pellet was finally resuspended in 150 µL of Neurobasal medium overnight at 4°C with slow rocking. Aliquots were frozen at -80°C.

### Methods

#### Syt1 knockdown, Abobotulinumtoxin A treatment and immunofluorescence staining of hippocampal neurons

For the assessment of BoNT/A toxic function following Syt1 KD with CRISPRi and shRNA, E18 hippocampal neurons were cultured on glass-bottom dishes (CellVis, Cat. #D29-10-1.5-N) at 50,000 neurons per 78.54 $mm^2$ confluency to ensure mature and functional synaptic connections. For CRISPRi Syt1 KD, neurons (DIV14) were transduced with dCas9-KRAB/TagBFP2 (control) or three different CRISPRi Syt1-targetting sgRNAs (sgRNA1-3) that were also designed to produce TagBFP2 marker, for 7 days. The lentiviral transduction of the neurons was performed as follows: 20 µL of the respective lentiviral stocks were diluted into 30 µL of conditioning medium collected from the dishes, with the remaining conditioning medium being collected into a sterile 15 mL Falcon tube. The virus mix was added onto the neurons, and incubated for 4-6 h at 37ºC in a 5% $CO_2$ atmosphere, after which the remaining conditioning medium was added back onto the dishes. For rescue experiments, the neurons were subsequently transduced with pLenti6.3-Syt1wt-pH on DIV19 for 72 h as described above. On DIV21-22, the conditioning medium was collected from the dishes, and the neurons were stimulated with high $K^+$ buffer supplemented with 10 U of Abobotulinumtoxin A (Ipsen; Dysport®; BoNT/A-based licensed drug) for 5 min at 37ºC in 5% $CO_2$ atmosphere, washed three times with low $K^+$ buffer, and chased for 30 min in conditioning medium at 37ºC in a 5% $CO_2$ atmosphere. Samples were then either fixed for immunofluorescence staining (or processed for patch-clamp electrophysiology, as described below). Immunostaining was performed as described above with the exception of using primary antibodies recognizing endogenous Syt1 (Abcam, Cat. #13259; 1:100 dilution in blocking buffer) and BoNT/A-cleaved

SNAP-25 (SNAP-25/A; a kind gift from D. Sesardic and P. Stickings, Division of Bacteriology, National Institute for Biological Standards and Control, United Kingdom; 1:200 dilution in blocking buffer), after which the samples were mounted, and imaged using a spinning-disk confocal system (Marianas; 3I, Inc.) consisting of an Axio Observer Z1 (Carl Zeiss) equipped with a CSU-W1 spinning-disk head (Yokogawa Corporation of America), ORCA-Flash4.0 v2 sCMOS camera (Hamamatsu Photonics), and 20x 0.8 NA / Plan-Apochromat / 550 µm WD objective. Image acquisition was performed using SlideBook 6.0 (3I, Inc). The CRISPRi results presented are from n=21 random regions of interest (ROIs) for both conditions, originating from two independent experiments, and the dot plots indicate the average fluorescence intensity±SEM of endogenous Syt1 and SNAP-25/A. The pLenti6.3-Syt1$^{wt}$-pH experiments are from n=3 random ROIs of both Syt1 sgRNA and control conditions (one independent experiment), and the dot plots indicate the average fluorescence intensity±SEM of endogenous Syt1 and cleaved SNAP-25.

As Syt1 is a long-lived protein that is challenging to KD (*14*), we used an shRNA KD strategy to achieve Syt1 KD in cultured hippocampal neurons (KD for 14 days). For shRNA-induced Syt1 KD, E18 hippocampal neurons (at DIV7) were co-transfected with EGFP and pRNAi-mU6-Green with sh-lacZ insertion (Biosettia, Cat. #SORT-A15) or Syt1 shRNA in a 1:1 ratio using Lipofectamine 2000. On DIV21-22, the conditioning medium was collected from the culture plates and the neurons were stimulated with high $K^+$ buffer supplemented with 10 U of Abobotulinumtoxin A for 5 min at 37ºC in a 5% $CO_2$ atmosphere, then washed three times with low $K^+$ buffer, after which the buffer was replaced with the conditioning medium, and the neurons were chased for 30 min or 16 h at 37ºC in a 5% $CO_2$ atmosphere. Samples were then fixed and processed for immunofluorescence staining and imaged with a spinning-disk confocal system as described above. The results presented for 30 min Abobotulinumtoxin A treatment are from n=26 (control) and n=27 (Syt1 shRNA) EGFP-positive neurons and for 16 h Abobotulinumtoxin A treatment are from n=29 (control) and n=30 (Syt1 shRNA) EGFP-positive neurons, from two independent experiments; the dot plots indicate the average of the fluorescence intensity±SEM of endogenous Syt1 and cleaved SNAP-25 from individual EGFP-positive somas.

**Electrophysiology**

To study the toxic function of Abobotulinumtoxin A following Syt1 KD, whole-cell patch-clamp electrophysiology experiments were performed on E18 hippocampal neurons at DIV21-22. CRISPRi sgRNA1 KD of Syt1 and 30 min Abobotulinumtoxin A treatment were done as described above. Experiments were performed at room temperature (20–23 ºC) using an Axopatch 700B amplifier (Molecular Devices) and Axograph software (Axograph Scientific). Cells were placed in a bath and continuously perfused with extracellular solution containing 140 mM NaCl, 5 mM KCl, 2 mM $CaCl_2$, 1 mM $MgCl_2$, 10 mM HEPES, and 10 mM D-glucose, adjusted to pH 7.4 with NaOH. Patch pipettes, fabricated from borosilicate glass capillaries (Harvard Apparatus), were pulled using a horizontal puller (Sutter Instruments), with a resistance of 3–6 MΩ, and fire-polished. The pipettes were filled with an intercellular solution containing 145 mM CsCl, 2 mM $CaCl_2$, 2 mM $MgCl_2$, 10 mM EGTA, and 2 mM MgATP, adjusted to pH 7.4 with CsOH. Spontaneous miniature inhibitory postsynaptic currents (mIPSCs) were recorded at a holding potential of −70 mV, in the presence of 1 nM tetrodotoxin (bath-applied; Abcam Cat. #120055) to block action potentials. Signals were filtered at 5 kHz and sampled at 20 kHz. Recordings with a series resistance above 20 MΩ were discarded. Peak amplitudes and frequencies were calculated using Axograph X (Axograph Scientific). Peak amplitudes were detected with a 3:1 signal to noise ratio as the threshold, and all peaks were manually examined to select well-separated events. Peak amplitudes calculated by Axograph X for each event were averaged to determine the final values for each cell. Frequency values were calculated by including all postsynaptic events above the 3:1 signal to noise ratio for a recording period of at least 10 min. The presented results are from n=5 (naïve), n=8 (control; dCas9-KRAB/TagBFP2), n=12 (control + 30 min Abobotulinumtoxin A; dCas9-KRAB/TagBFP2 following 30 min Abobotulinumtoxin A-treatment), n=9 (Syt1 sgRNA1), and n=11 (Syt1 sgRNA1 + 30 min Abobotulinumtoxin A treatment) recordings from 2 independent experiments. The dot plots indicate the average from individual recordings ± SD.

**HRP cytochemistry and electron microscopy**

Electron microscopy (EM) analysis was performed on cultured E18 hippocampal neurons grown on dishes (CellVis, Cat. #D29-10-1.5-N) at 50,000 neurons per 78.54 $mm^2$ confluency to ensure mature and functional synaptic connections. On DIV21-22, the neurons were stimulated with high $K^+$ buffer supplemented with 5 µg $mL^{-1}$ BoNT/$Ai^{wt}$-HRP, BoNT/$Ai^{PSG}$-HRP BoNT/$Ai^{SV2}$-HRP and BoNT/$Ai^{PSG,SV2}$-HRP for 5 min at 37ºC in a 5% $CO_2$ atmosphere, washed with low $K^+$ buffer (8-10 mL) and chased for 10 min at 37ºC in a 5% $CO_2$ atmosphere. They were then fixed with 2% paraformaldehyde, 1.5% glutaraldehyde (Electron Microscopy Sciences, Cat. #16210) in 0.1 M sodium cacodylate (Sigma-Aldrich, Cat. #C0250) buffer, pH 7.4, for 20 min at room temperature, washed three times for 3 min with 0.1 M sodium cacodylate buffer, and processed for 3,3′-diaminobenzidine tetrahydrochloride (DAB; Sigma-Aldrich Cat. #D5905) cytochemistry using standard protocols. Samples were contrasted with 1% osmium tetroxide and 2% uranyl acetate before dehydration and embedded in LX-112 or EPON resin using a BioWave tissue processing system (Pelco) as previously described (*15*). Thin sections (80-90 nm) were cut using an ultramicrotome (Leica Biosystems, UC6FCS), and imaged with a transmission electron microscope (JEOL USA, Inc. model 1101) equipped with a cooled charge-coupled device camera (Olympus; Morada CCD Camera).

Images were acquired when HRP precipitate was observed. For the quantification of the percentages (%) of total HRP signal, the localization of the HRP-tagged holotoxins was quantified manually based on the following criteria: i) endocytic structures smaller than 80 nm in diameter were assigned as synaptic vesicles, ii) endocytic structures larger than 80 nm in diameter with round morphology were assigned as endosomes, and structures of similar diameter and concave appearance or hollow center were assigned as early endosomes (all counted together as endosomes), iii) large endocytic structures with a multivesicular appearance were assigned as multivesicular bodies, iv) large double-membrane endocytic structures that contained cellular material were assigned as autophagosome, as we have described earlier (*16, 17*), v) tubular structures approximately 50-80 nm in diameter, and free of ribosomes, were assigned as tubular structures, and vi) separate HRP precipitates connected to the extracellular side of the plasma membrane were assigned as plasma membrane staining. Interconnected tubular structures with more than one HRP precipitate were counted as one structure. In total, 376, 308, 295 and 206 structures containing HRP precipitate, or separate HRP precipitates on the plasma membrane, were counted for BoNT/$Ai^{wt}$-HRP, BoNT/$Ai^{PSG}$-HRP, BoNT/$Ai^{SV2}$-HRP and BoNT/$Ai^{PSG,SV2}$-HRP, respectively. For the CRISPRi experiments, 394 and 464 structures containing HRP precipitate, or separate HRP precipitates on the plasma membrane, were counted for BoNT/$Ai^{wt}$-HRP in dCas9-KRAB/TagBFP2 (control) and CRISPRi sgRNA1 Syt1 KD neurons, respectively. The results are presented as average ± SEM. For the quantification of average sectional area ($\mu m^2$) of the endocytic profiles containing the HRP precipitate of a given holotoxin, individual HRP-stained profiles were segmented and the area was measured in ImageJ/Fiji (*18, 19*)(

https://imagej.nih.gov/ij/), with the dot plots indicating the average area (μ$m^2$) of an endocytic profile ± SEM.

For Ruthenium red (RuR; Sigma-Aldrich, Cat. #00541) staining, E18 hippocampal neurons were lentivirally transduced on DIV14 with dCas9-KRAB/TagBFP2 (control) and CRISPRi Syt1 sgRNA1 KD for 7 days as described above. On DIV22, neurons were stimulated with high $K^+$ buffer for 5 min at 37ºC in a 5% $CO_2$ atmosphere, washed with low $K^+$ buffer (8-10 mL) and then chased for 10 min at 37ºC in a 5% $CO_2$ atmosphere. They were then briefly rinsed with 0.1 M sodium cacodylate buffer pH 7.4 and fixed with 2.5% glutaraldehyde in 0.1 M sodium cacodylate buffer pH 7.4 supplemented with 1 mg $mL^{-1}$ RuR for 1 h at room temperature, washed three times for 10 min with 0.1 M sodium cacodylate buffer, and treated with 1% osmium tetroxide in 0.1 M sodium cacodylate buffer supplemented with 1 mg $mL^{-1}$ RuR for 3 h at room temperature. Following 3 washes with 0.1 M sodium cacodylate buffer, the samples were dehydrated and embedded in EPON resin using a BioWave tissue processing system (Pelco) as previously described (*15*), thin sectioned (80-90 nm sections) and imaged as above.

**Electron tomography**

E18 rat hippocampal neurons were induced on DIV14 with CRISPRi Syt1 sgRNA1 KD for 7 days and processed for EM as described above. For Syt1 KD electron tomography, EM samples were prepared as above. Approximately 200 nm resin sections were then cut on a Leica Ultracut 6 ultramicrotome, and the grid was subsequently coated with a thin carbon layer. Tomography was completed as described previously (*20*). In brief, a dual-axis tilt series spanning ±60° with 1° increments were acquired on a Tecnai F30 transmission electron microscope (FEI) at 300 kV using a Gatan one view camera under the control of Serial EM. Tilt series were reconstructed using weighted back-projection and patch tracking in IMOD (https://bio3d.colorado.edu/imod/). Segmentation was performed by density thresholding using the Isosurface Render program in IMOD for specific ROIs.

**Mouse phrenic nerve hemidiaphragm assay**

To demonstrate that the At647N labeling of BoNT/A holotoxin did not impair its mechanism of action, the potency of wild-type active BoNT/A-At647N was determined in comparison to non-labeled wild-type active BoNT/A using the mouse phrenic nerve hemidiaphragm (MPN) assay. The MPN assay was performed using 20–30 g Swiss mice (Janvier SA, France) as described previously (*21*). According to §4 Abs. 3 (sacrificing animals for scientific purposes, German animal protection law (TSchG)), the number of animals sacrificed by trained personnel before dissection of organs was reported yearly to the animal welfare officer of the Central Animal Laboratory and to the local authority, Veterinäramt Hannover. Isolated *N. phrenicus* hemidiaphragm tissue was then treated with non-labeled or BoNT/A-At647N, inducing a characteristic time-dependent decrease of the contraction amplitude of the indirectly stimulated muscle (*21*). Employing the dose-response curve previously established for recombinant BoNT/A (*11*), catalytically active BoNT/A-At647N retained high potency (~20%) of the non-labelled BoNT/A. This reduction is acceptable considering the up to one order of magnitude lot-to-lot variation of specific toxicity of BoNT/A isolated from *C. botulinum* cultures. We estimate that the impairment to the mechanism of action caused by the At647N labeling of BoNT/Ai$^{wt}$, BoNT/Ai$^{PSG,}$ BoNT/Ai$^{SV2}$ and BoNT/Ai$^{PSG,SV2}$ is highly likely to be in a similar range to that of BoNT/A-At647N given that all single-site mutations (receptor binding site mutations

W1266L/G1141D/G1292R, and LC inactivation E224A/R363A/Y366F) neither remove nor introduce the lysine residues that the NHS-ester targets.

**uPAINT imaging**

Universal point accumulation imaging in nanoscale topography (uPAINT) experiments were performed as described earlier (*12, 22*). For imaging the single-molecule mobility of BoNT/Ai$^{wt}$-At647N, BoNT/Ai$^{PSG}$-At647N, BoNT/Ai$^{SV2}$-At647N and BoNT/Ai$^{PSG,SV2}$-At647N holotoxins on the plasma membrane, E18 hippocampal neurons were grown on super-resolution-compatible glass-bottom dishes (*1*), and on DIV21-22 the growth medium was replaced with low $K^+$ buffer (0.5 mM $MgCl_2$ (Chem-Supply, Cat. #MA029), 2.2 mM $CaCl_2$ (Sigma-Aldrich, Cat. #C5080), 5.6 mM KCl (Ajax Finechem Pty Limited, Cat. #1206119), 145 mM NaCl, 5.6 mM D-glucose (AMRESCO, Cat. #0188), 0.5 mM ascorbic acid (Sigma-Aldrich, Cat. #A5960), 0.1% (wt/vol) bovine serum albumin (BSA; Sigma-Aldrich, Cat. #A8022) and 15 mM HEPES (Sigma-Aldrich, Cat. #H3375), pH 7.4, 290-310 mOsm). Neurons were then transferred to the microscope, and stimulated with high $K^+$ buffer (56 mM KCl, 0.5 mM ascorbic acid, 0.1% BSA, 15 mM HEPES, 5.6 mM D-glucose, 95 mM NaCl, 0.5 mM $MgCl_2$, and 2.2 mM $CaCl_2$, at pH 7.4, 290–310 mOsm) supplemented with 100 pM holotoxins using a custom-made perfusion system described earlier (*1*). The single-molecule mobility of the holotoxins landing on the plasma membrane was then immediately imaged using a Roper Scientific iLas$^2$ Ring-TIRF (total internal reflection fluorescence) microscope with a CFI Apo 100×/1.49 N.A. an oil-immersion objective (Nikon Instruments), an Evolve 512 Delta EMCCD camera (Photometrics) mounted on a TwinCam LS Image Splitter (Cairn Research), a Perfect Focus System (Nikon), an iLas$^2$ double-laser illuminator (Roper Scientific) for 360° TIRF illumination, and a 642 nm laser (100 mW, Vortran). TetraSpeck microspheres (ThermoFisher Scientific, Cat. #T7279) were used for TIRF angle calibration prior to imaging. Image acquisition was performed using MetaMorph software (Molecular Devices, Version 7.10.2). The uPAINT results for BoNT/Ai$^{wt}$-At647N, BoNT/Ai$^{PSG}$-At647N, BoNT/Ai$^{SV2}$-At647N and BoNT/Ai$^{PSG,SV2}$-At647N are shown from n=17 acquisitions from randomly chosen ROIs (i.e. number of individual neuronal cultures; one acquisition per culture), originating from 5-6 independent experiments (i.e. independent embryonic recoveries). The average number of tracks ±SEM per acquisition was 5800±1100 for BoNT/Ai$^{wt}$-At647N, 1800±300 for BoNT/Ai$^{PSG}$-At647N, 4200±1100 for BoNT/Ai$^{SV2}$-At647N and 2400±400 for BoNT/Ai$^{PSG,SV2}$-At647N, with the dot plots indicating the average mobility±SEM in each acquisition.

For imaging SV2A and Syt1 mobility on the plasma membrane and to address the receptor mobility changes upon exposure to respective BoNT/Ai toxins, single channel uPAINT (for all control experiments without addition of toxin) and dual-color uPAINT (for all experiments on the effects of toxins on receptor mobility; see further details on the dual-color uPAINT imaging below) were carried out separately. To image SV2A$^{wt}$-pHluorin (*23*) (SV2A$^{wt}$-pH) and SV2A$^{T84A}$-pHluorin (*23*) (SV2A$^{T84A}$-pH) single-molecule mobility on the plasma membrane in the absence of toxins, uPAINT experiments were carried out as described previously (*12*) and above, with the following exceptions. Neurons were transfected with SV2A$^{wt}$-pH or SV2A$^{T84A}$-pH on DIV14. On DIV21-22, the neurons were stimulated with high $K^+$ buffer supplemented with 100 pM anti-GFP Atto565 nanobodies (At565nb; anti-GFP sdAb antibody - FluoTag-Q At565; Synaptic Systems Cat. #N0301-At565-S). High $K^+$ stimulation increases the extracellular exposure of the SV2 luminal epitope following synaptic vesicle fusion with the plasma membrane (*24-28*). At565nb specifically bind to the intravesicular pHluorin-tag of the constructs following synaptic vesicle fusion with the plasma membrane and exposure of the pHluorin-tag to the extracellular space. The mobility of SV2A$^{wt}$-pH/At565nb and SV2A$^{T84A}$-pH/At565nb was then immediately recorded using TIRF imaging. The ROIs for imaging were chosen based on the pHluorin fluorescence signal of the transfected neurons and observed neuronal morphology. The uPAINT high $K^+$ stimulation results for SV2A$^{wt}$-pH/At565nb with and without BoNT/Ai$^{wt}$-At647N exposure are shown from n=23 acquisitions and n=11 acquisitions for SV2A$^{wt}$-pH/At565nb with and without BoNT/Ai$^{SV2}$-At647N exposure from 3-5 independent experiments. The uPAINT high $K^+$ stimulation results for SV2A$^{T84A}$-pH/At565nb are shown from n=17 acquisitions for both conditions from 3 independent experiments. The number of acquisitions refers to individual neuronal cultures (one acquisition per culture) and the number of independent experiments refers to independent embryonic recoveries. The average number of tracks±SEM per acquisition was 1000±100 for SV2A$^{wt}$-pH/At565nb (control for SV2A$^{wt}$-pH/At565nb+BoNT/Ai$^{wt}$-At647N), 1000±200 for SV2A$^{wt}$-pH/At565nb (control for SV2A$^{wt}$-pH/At565nb+BoNT/Ai$^{SV2}$-At647N), and 900±200 for SV2A$^{T84A}$-pH/At565nb (control for SV2A$^{T84A}$-pH/At565nb+BoNT/Ai$^{wt}$-At647N), and the dot plots indicate the average mobility±SEM from each acquisition.

To image Syt1$^{wt}$-pHluorin (*29, 30*) (Syt1$^{wt}$-pH; A kind gift from Prof. Volker Haucke, The Leibniz-Institute for Molecular Pharmacology), Syt1$^{K52A}$-pHluorin (Syt1$^{K52A}$-pH) and Syt1$^{K326A,K328A}$-pHluorin (*23*) (Syt1$^{K326A,K328A}$-pH) single-molecule mobility on the plasma membrane in the absence of holotoxins, uPAINT experiments were carried out as described above by transfecting the neurons with the receptive constructs. The ROIs for imaging were chosen based on the pHluorin fluorescence signal of the transfected neurons and observed neuronal morphology. The uPAINT high $K^+$ stimulation (control) results for Syt1$^{wt}$-pH/At565nb, Syt1$^{K52A}$-pH/At565nb and Syt1$^{K326A,K328A}$-pH/At565nb are shown from n=14 acquisitions/condition originating from 3-4 independent experiments. The number of acquisitions refers to individual neuronal cultures (one acquisition per culture) and the number of independent experiments refers to independent embryonic recoveries. The average number of tracks ±SEM per acquisition were 8900±2200 for Syt1$^{wt}$-pH/At565nb (control for Syt1$^{wt}$-pH/At565nb+BoNT/Ai$^{wt}$-At647N), 5900±1000 for Syt1$^{K52A}$-pH/At565nb (control for Syt1$^{K52A}$-pH/At565nb+BoNT/Ai$^{wt}$-At647N), and 4900±800 SEM for Syt1$^{K326A,K328A}$-pH/At565nb (control for Syt1$^{K326A,K328A}$-pH/At565nb+BoNT/Ai$^{wt}$-At647N), and the dot plots indicate the average mobility ± SEM in each acquisition.

**Dual-color uPAINT imaging**

The mobilities of SV2A$^{wt}$-pH/At565nb and SV2A$^{T84A}$-pH/At565nb following high K$^{+}$ stimulation supplemented with 100 pM BoNT/Ai$^{wt}$-At647N or BoNT/Ai$^{SV2}$-At647N were acquired simultaneously with dual-color uPAINT imaging, as described earlier (*31*), and the receptor mobility was analyzed separately. In short, E18 hippocampal neurons were grown on super-resolution-compatible glass-bottom dishes (*1*) and, on DIV14, transfected with SV2A$^{wt}$-pH or SV2A$^{T84A}$-pH. On DIV21-22, uPAINT imaging was carried out by stimulating the neurons with high K$^{+}$ buffer supplemented with 100 pM of BoNT/Ai$^{wt}$-At647N or 100 pM BoNT/Ai$^{SV2}$-At647N, and 100 pM anti-GFP At565nb, and the mobilities of the At647N-labeled holotoxin and SV2A$^{wt}$-pH/At565nb and SV2A$^{T84A}$-pH/At565nb were simultaneously recorded using TIRF microscopy. We used the imaging setup described above with the following additions. Two Evolve 512 Delta EMCCD cameras (Photometrics) were employed to record simultaneously on two channels, with a TIRF-quality ultra-flat quadruple beam splitter (ZT405/488/561/647rpc; Chroma Technology) for distortion-free reflection of lasers and QUAD emission filter (simultaneous imaging quadruple laser filter set; ZET405/488/561/640 m; Chroma) for dual imaging. TetraSpeck microspheres were used for dual-camera alignment and TIRF angle calibration prior to imaging.

The uPAINT results for SV2A$^{wt}$-pH/At565nb and SV2A$^{T84A}$-pH/At565nb following holotoxin treatment are shown from n=11-23 acquisitions per condition from 3-5 independent experiments. The average number of tracks±SEM per acquisition was 1100±150 for SV2A$^{wt}$-pH/At565nb (following BoNT/Ai$^{wt}$-At647N exposure) and 900 ± 400 SEM for SV2A$^{T84A}$-pH/At565nb (following BoNT/Ai$^{wt}$-At647N exposure), and the dot plots indicate the average mobility±SEM in each acquisition.

Syt1$^{wt}$-pH/At565nb, Syt1$^{K52A}$-pH/At565nb and Syt1$^{K326A,K328A}$-pH/At565nb mobility data following 100 pM BoNT/Ai$^{wt}$-At647N-treatment were acquired with a similar dual-color uPAINT imaging setup. The number of acquisitions refers to individual neuronal cultures (one acquisition per culture). ROIs were chosen based on the transfected neuron pHluorin fluorescence and neuronal morphology. The average number of tracks±SEM per acquisition were 7700±2900 for Syt1$^{wt}$-pH/At565nb, 3500±600 for Syt1$^{K52A}$-pH/At565nb, and 2800±500 for Syt1$^{K326A,K328A}$-pH/At565nb, all following BoNT/Ai$^{wt}$-At647N exposure, and the dot plots indicate the average mobility ± SEM in each acquisition.

**sdTIM imaging**

Subdiffractional tracking of internalized molecules (sdTIM) experiments were performed as described earlier (*1, 12*). To image the single-molecule mobility of BoNT/Ai$^{wt}$-At647N, BoNT/Ai$^{PSG}$-At647N, BoNT/Ai$^{SV2}$-At647N or BoNT/Ai$^{PSG,SV2}$-At647N holotoxins following internalization, E18 hippocampal neurons were grown on super-resolution-compatible glass-bottom dishes, and on DIV21-22 the growth medium was replaced with low $K^+$ buffer. The neurons were then stimulated with high $K^+$ buffer supplemented with 1 nM BoNT/Ai$^{wt}$-At647N, BoNT/Ai$^{PSG}$-At647N, BoNT/Ai$^{SV2}$-At647N or BoNT/Ai$^{PSG,SV2}$-At647N holotoxins for 5 min at

37°C in a 5% $CO_2$ atmosphere, washed three times with low $K^+$ buffer (8-10 mL) to remove unbound toxins and chased for 10 min at 37°C in a 5% $CO_2$ atmosphere to induce endocytic uptake of the holotoxins. They were then imaged as described above at 50 Hz (16,000 frames by image streaming) and 20 ms exposure, at 37°C in HILO (highly inclined and laminated optical sheet) illumination (*32*). TetraSpeck microspheres were used for dual-camera alignment and HILO angle calibration prior to imaging. sdTIM results are shown from n=17 acquisition per condition from randomly chosen ROIs, originating from 5-6 independent experiments (number of independent embryonic recoveries). The number of acquisitions refers to individual neuronal cultures (one acquisition per culture). The average number of tracks ± SEM per acquisition was 2100±700 for BoNT/Ai$^{wt}$-At647N, 400±50 for BoNT/Ai$^{PSG}$-At647N, 800±200 for BoNT/Ai$^{SV2}$-At647N and 500±80 for BoNT/Ai$^{PSG,SV2}$-At647N, and the dot plots indicate the average mobility±SEM.

**Retrograde transport assay**

Hippocampal neurons were collected from E18 rats and seeded in the poly-L-lysine-coated chambers of microfluidic devices as previously described (*1*) at 50,000 and 5,000 neuron confluency in the soma and nerve terminal chambers, respectively. Using microfluidic devices enables neuronal culturing in an orientation where the neuronal soma, axons and the nerve terminals are separated in specific compartments. A pulse-chase assay with holotoxins was carried out as described previously(*17*) at DIV21. Briefly, the neuronal conditioning medium from both chambers was collected and the soma chamber was stimulated with high $K^+$ buffer and the nerve terminal chamber with high $K^+$ buffer supplemented with 100 nM BoNT/A$^{wt}$-At647N, BoNT/A$^{PSG}$-At647N, BoNT/A$^{SV2}$-At647N or BoNT/A$^{PSG,SV2}$-At647N for 5 min at 37ºC in a 5% $CO_2$ atmosphere. Both chambers were then washed with low $K^+$ buffer to remove unbound toxins, and neurons were chased for 2 h in the collected conditioning medium at 37ºC in a 5% $CO_2$ atmosphere. The retrograde transport carriers containing the fluorescently labeled toxins in the axons were recorded using a spinning-disk confocal microscope with the following settings: a 60x oil objective was used with a 1.4 NA / 130 µm WD / 91.5 nm/pixel to obtain close to Nyquist (57 nm/pixel) condition images, with a pinhole aperture of 100 nm. An AndorZyla sCMOS camera was used with a time loop set at 300 ms for 90 s. Three-hundred-frame videos were deconvolved with Huygens Professional 19.10 software (Scientific Volume Imaging; https://svi.nl/Huygens-Professional) and retrograde carriers were then tracked using the Simple Linear Assignment Problem (LAP) Tracker of the TrackMate(*33*)(https://imagej.net/plugins/trackmate/) plugin for ImageJ with the following options: linking max distance: 5 µm; gap-closing max distance: 5 µm; gap-closing maximum frame gap: 2. Tracks shorter than 9 s (10% total time) were excluded. Kymographs were generated using the Multi-Kymograph plugin for ImageJ. The results presented are from n=28-31 channels/condition from 4 independent experiments, and the dot plots indicate the average frequency±SEM from an individual axonal channel.

**Single-molecule and retrograde transport carrier tracking**

The single-molecule localization and dynamics were extracted from the 16,000 frame TIRF and HILO acquisitions as described previously (*12, 34*). BoNT/Ai$^{wt}$-At647N, BoNT/Ai$^{PSG}$-At647N, BoNT/Ai$^{SV2}$-At647N and BoNT/Ai$^{PSG,SV2}$-At647N, as well as SV2A$^{wt}$-pH/At565nb, SV2A$^{T84A}$-pH/At565nb, Syt1$^{wt}$-pH/At565nb, Syt1$^{K52A}$-pH/At565nb and Syt1$^{K326A,K328A}$-pH/At565nb, were detected and tracked using a combination of wavelet segmentation (*35*) and simulated annealing (*36*). PALMTracer (*34, 37*) in Metamorph software (MetaMorph Microscopy Automation and Image Analysis Software, v7.7.8; Molecular Devices) was used to obtain the mean square displacement (MSD) and diffusion coefficient (D; $\mu m^2 s^{-1}$) values. Tracks shorter than eight frames were excluded from the analysis to minimize nonspecific background. Cross-correlation drift correction of the uPAINT data was done using the SharpViSu tool (*38*) (https://github.com/andronovl/SharpViSu) in MATLAB 2017b (https://au.mathworks.com/matlabcentral/answers/498405-matlab-2017b-download). The $Log_{10}D$ immobile and mobile fraction distributions were calculated as previously described(*39*), setting the displacement threshold to 0.03 $\mu m^2 s^{-1}$ (*i.e.* dotted line in the graph $Log_{10}D = -1.45$ when [D] = $\mu m^2 s^{-1}$ as described previously (*12, 39*)). The mobile to immobile (M/IMM) ratio was determined based on the frequency distribution of the diffusion coefficients ($Log_{10}D$) of immobile ($Log_{10}D \leq -1.45$) and mobile ($Log_{10}D > -1.45$) molecules (the immobile fraction of molecules represents BoNT/$A^{wt}$-At647N molecules for which the displacement within 4 frames was below the spatial detection limit of our methods, 106 nm). The area under the MSD curve (AUC) was calculated in Prism 9 for macOS version 9.1.1 (GraphPad Prism 9 for macOS; https://www.graphpad.com/scientific-software/prism/). The super-resolved image color-coding was done as previously described (*1*) using ImageJ/Fiji (2.0.0-rc-43/1.50e; National Institutes of Health), with each colored pixel in the average intensity maps indicating the localization of an individual molecule (bar: 8 to 0, high to low density), the color-coded pixels in the average diffusion coefficient map presenting an average value for each single-molecule track at the site of localization (bar: $Log_{10}$ 1 to -5, high to low mobility), and the color-coding of the track maps representing the detection time (bar: 0-16,000 frame acquisition) point during acquisition.

***In situ* proximity ligation assay**

Hippocampal neurons from E18 rats were seeded on glass-bottom dishes (CellVis, Cat. #D29-10-1.5-N) at 50,000 neurons per 78.54 $mm^2$ confluency to ensure mature and functional synaptic connections. On DIV21-22, neurons were left untreated (naïve control), stimulated for 5 min in high $K^+$ buffer (vehicle control), or stimulated for 5 min in high $K^+$ buffer supplemented with 100 pM BoNT/$Ai^{wt}$-At647N, BoNT/$Ai^{GT1b}$-At647N, BoNT/$Ai^{SV2}$-At647N or BoNT/$A^{PSG,SV2}$-At647N. Neurons were then fixed with 4% paraformaldehyde in PBS for 20 min at room temperature, washed three times with PBS, permeabilized with 0.1% Triton X-100 for 4 min, washed three times with PBS, and processed for the *in situ* proximity ligation assay (PLA) according to the manufacturer's instructions (DuoLink, Merck, Cat. #DUO92101). Every experiment was performed with a pair of primary antibodies against endogenous Syt1 (Anti-Syt1, Abcam, Cat. #ab13259; 1:200 dilution) and SV2C (Anti-SV2C, Synaptic Systems, Cat. #119203; 1:200 dilution). Neurons were mounted in Duolink *in situ* mounting medium containing DAPI. Optical sections of the fluorescent signal of PLA dots and DAPI, spanning entire neurons, were acquired from ten random ROIs in each sample. Imaging was done using a spinning-disk Marianas 3I, Inc confocal system as described earlier, with the exception of using a 63x 1.4 NA/Plan-Apochromat/180 µm WD objective. For image analysis, a 200x200 pixel area was selected manually around each nucleus (based on DAPI labeling, with overlapping nuclei being excluded) and the mid-section of the nucleus was determined based on the peak fluorescence intensity of DAPI using the Plot z-axis Profile-function in ImageJ/Fiji. To quantify Syt1-SV2C interactions at the synaptic connections in the somatodendritic area, the site of synaptic contacts, and to avoid the uniform non-specific PLA signal observed on the glass bottom of the dish, a maximum intensity projection of the PLA signal was then obtained from ± 10 optical slice (z-step 0.2 µm) offsetting from the nuclear midsection of each neuron. Automated image analysis to identify and quantify the number of PLA dots (reflecting the detection of a Syt1-SV2C protein-protein interaction) using ImageJ/Fiji (Version 2.1.0/1.53c) was then conducted as described previously (*40*). The results presented are from n=25 randomly chosen 200x200 pixel ROIs encompassing the DAPI staining, from two independent experiments for each condition, and the dot plots indicate the number of PLA dots per nucleus±SEM (with neurons originating from embryos of different dams).

**Fluorescence emission assay in fixed neurons**

For fluorescence emission step analysis (fig. S7M-S7O) of endocytosed BoNT/Ai$^{wt}$-At647N, BoNT/Ai$^{PSG}$-At467N, BoNT/Ai$^{SV2}$-At467N, and BoNT/Ai$^{PSG,SV2}$-At467, E18 hippocampal neurons were transfected with pEGFP-C1 (BD Biosciences Clontech, Cat. #6084-1) on DIV14, and subjected to sdTIM on DIV21-22 by inducing the activity-dependent uptake of the indicated holotoxins as described above. Neurons were then fixed with 4% paraformaldehyde (Electron Microscopy Sciences, Cat. #15710) in PBS for 30 min at room temperature, followed by blocking with 1% BSA (Sigma-Aldrich, Cat. #A8022) in PBS for 30 min and permeabilization with 0.05% Triton X-100 (Sigma-Aldrich, Cat. #T-9284) in PBS for 4 min at room temperature. Samples were then incubated for 1 h at room temperature with primary antibody against synapsin-1 (Synaptic Systems, Cat. #106011, 1:200 dilution in blocking buffer). After three 5 min washing steps in PBS, the samples were incubated with Alexa Fluor 546 goat anti-mouse IgG (Invitrogen, Cat. #A11030, 1:1000 dilution in blocking buffer) secondary antibody for 30 min at room temperature shielded from light; after three washes in PBS, they were mounted in ProLong Gold Antifade Mountant (ThermoFisher Scientific, Cat. #P10144). The fluorescence emission of BoNT/A$^{wt}$-At647N and mutants from the fixed neurons was recorded using a Roper iLas$^2$ TIRF microscope at 50 Hz over 16,000 frames (20 ms exposure time), similar to the sdTIM live-cell imaging described above. The fluorescence emission steps of the toxins in presynaptic areas (positive for synapsin-1) were quantified by segmenting presynaptic ROIs based on overlapping EGFP (axons) and synapsin-1 (presynapses) signal, plotting the fluorescence emission curve in ImageJ/Fiji, and counting the number of resulting emission steps manually. The scatter plot shows the number of fluorescence emission steps of indicated holotoxins per presynapse, normalized to the DoL of each toxin. The results shown are from n=71-116 presynaptic ROIs per condition originating from 3 independent experiments, and the dot plots indicate the average fluorescence intensity per presynapses ± SEM.

**Automated image segmentation and fluorescence intensity analysis**

The quantification of Syt1 and cleaved SNAP-25 fluorescence following CRISPRi (Figure 5A-C) was done using CellProfiler 3(*41*) (https://cellprofiler.org/) and ImageJ/Fiji. Using ImageJ/Fiji, confocal stacks of the Abobotulinumtoxin A-treated and immunostained hippocampal neurons were Z-projected using the sum of fluorescence. The 2D Z-projections were then used in CellProfiler 3 to determine the area positive for TagBFP2 (a marker to identify neurons that had received the dCas9-KRAB and the respective sgRNA). First, the brightest TagBFP2 fluorescent spots were identified using the Identify Primary Object tool of CellProfiler 3 and the otzu algorithm (green areas highlighted in fig. S1A). Next, these primary spots were expanded following the TagBFP2 fluorescence signal until the whole TagBFP2 ROI of each image was segmented (blue areas highlighted in fig. S1A). The mean fluorescence intensity (MFI, in arbitrary units) of endogenous Syt1 and cleaved SNAP-25 was then measured from the segmented ROI. The mean background fluorescence intensity was determined manually from 15 areas outside the cells in each image. This value was subtracted from the MFI value, and the presented graphs show MFI ± SEM.

The MFI quantification for Syt1 shRNA knockdown (fig. S1B-S1H) was performed similar to the approach described above for CRISPRi by first Z-projecting the sum fluorescence of the 3D-stacks acquired with a spinning-disk confocal microscope as above. The 2D Z-projections were then used in CellProfiler 3 to segment the area positive for EGFP (a marker to identify neurons that had received the co-transfected EGFP with the respective shRNA). The values of threshold fluorescence intensity were chosen in CellProfiler 3 so that only the brightest signal of the EGFP-positive cells was automatically detected in all images. These bright areas corresponded to the neuronal soma (boxed area shown in higher magnification on right). The MFI of Syt1 and cleaved SNAP-25 was then quantified in these selected areas. SNAP-25 localization in differentiated neurons has been shown to be plasma membrane-associated in axonal compartments and not restricted to synaptic junctions, and also to localize in tubulo-vesicular structures in the cytoplasm of the soma and the axon, but rarely in the smaller dendrites (*42*). In the shRNA experiments, the Syt1 and SNAP-25/A MFI was quantified in the soma compartment, which is the site for many synaptic connections, similar to quantifications done in motorneurons previously (*43*), due to low neuronal transfections efficiency. In contrast, in the CRISPRi experiments above the MFI was quantified in whole neurons.

For pLenti6.3-Syt1$^{wt}$-pH recue of Syt1 sgRNA1 CRISPRi, 3D images acquired by spinning-disk confocal microscopy were Z-projected into 2D images using the 'Z-project sum of fluorescence intensities' function of the ImageJ. CellProfiler 3 was used to automatically identify TagBFP2-positive cells using a manually determined threshold of fluorescence estimated as > 2x the median background fluorescence outside the cells. The identified TagBFP2-positive areas corresponded to the soma of each neuron (the less bright dendrites and axons where not selected by the threshold of fluorescence). Within these TagBFP2-positive areas, the software measured the MFI of Syt1$^{wt}$-

pH, Syt1 (immunofluorescence of Syt1), and SNAP25/A (immunofluorescence staining of cleaved SNAP-25). Using the same software, from each 2D image, the median fluorescence of fifteen 20x20 pixel ROIs outlining individual neurons was manually determined for each of the four fluorescence channels, and these intensity values were subtracted from all the final measurements of each respective channel. The pLenti6.3-Syt1$^{wt}$-pH data are reported from n=3 random ROIs of both Syt1 sgRNA and control conditions (one independent experiment), and the dot plots indicate the MFI ± SEM of endogenous Syt1 and cleaved SNAP-25. The regression graphs (± 95% confidence intervals) are from representative Z-projected stacks of Syt1 sgRNA1 CRISPRi rescued with pLenti6.3-Syt1$^{wt}$-pH (Fig. 1G) and non-induced control neurons with pLenti6.3-Syt1$^{wt}$-pH (Fig. 1J). Graphs were made in Prism 9, and the $R^2$ for the regression analysis was determined in Excel (Microsoft Excel for Mac, version 16.55).

**Nanoscale spatiotemporal indexing clustering (NASTIC) analysis**

The NASTIC tool (*44*) was used to perform nanocluster analysis on the single-molecule localization data obtained with uPAINT. NASTIC utilizes the R-tree spatial indexing algorithm (*45*) to determine the overlap of the bounding boxes of individual molecular trajectories, as a measure of their potential membership in nanoclusters. The spatial extent of a cluster represents the convex hull encompassing the individual detections of all overlapping trajectories in the cluster. Extending the spatial indexing into the time dimension allowed us to expand the spatial nanoclustering into spatiotemporal clustering, such that trajectory bounding boxes must overlap in both space and time to be considered a cluster. This approach enabled us to determine clustering metrics such as apparent cluster lifetime and rate of formation (not shown), as well as identifying spatial "hotspots" of repeated cluster formation over the acquisition period. NASTIC employs several parameters: the radius factor (r) represents the multiplication of the radius of the ideal circularized extent of each trajectory used to construct the regular 2D spatial bounding region, and the time window (t) represents the temporal "thickness" of the resulting 3D bounding box. For these analyses, the default values of r = 1.2 and $t$ = 20 ms were used. Overlapping trajectories within ± 10 s in the same ROIs were counted as clusters. Clusters with a radius of > 150 nm were excluded from subsequent metrics. For visualization, identified clusters were colored according to the average detection time in the analysis, such that blue/red/green clusters represent those from early/middle/later in the analysis respectively. Hotspots of repeated cluster formation are highlighted in white. 2D kernel density estimations were performed using individual molecular detections, and were colored qualitatively such that higher density regions were represented with higher color temperature. Instantaneous diffusion coefficient plots of trajectories are colored such that smaller values are represented by higher color temperature (color bar range -5 to 1, $\log_{10}D$). Outliers were removed using the ROUT method with Q=1% in Prism 9. n=11-19 neurons/ condition from 3-5 independent experiments from the respective single-molecule experiments.

**Structural modeling of the SV2A-Syt1-GT1b-BoNT/A complex**

A structural model of the BoNT/A holotoxin in complex with tripartite GT1b-Syt1-SV2A, was compiled step-wise beginning with the full-length BoNT/A crystal structure (PDB 2NYY (*46*)) and full-length models of human SV2A and Syt1 from the AlphaFold2 database (https://alphafold.ebi.ac.uk (*47, 48*); https://github.com/deepmind/alphafold). The full-length SV2A and BoNT/A proteins were aligned with the co-crystal structure of the SV2C extracellular domain in complex with the 50 kDa BoNT/A $H_C$ fragment (PDB 4JRA (*49*)). The full-length Syt1 was then aligned to both the co-crystal structure of the GT1b ganglioside bound to the 50 kDa

BoNT/A $H_C$ fragment (*50*), and the co-crystal structure of the Syt1 C2B domain bound to the peptide of SV2A phosphorylated at Thr84 (*23*). The approximate alignment of Syt1 bound to GT1b was made based on the previously published molecular dynamics models of this complex (*51*). Alignments were performed in Pymol (Schrodinger), with adjustments to flexible regions in SV2A and Syt1 done manually to allow for their docking to bound partners in a physically reasonable way. Finally, bond lengths and angles were optimized in Coot (*52*)(https://www2.mrc-lmb.cam.ac.uk/personal/ pemsley/coot/). Structural images were made in Pymol (Schrodinger, USA; https://pymol.org/2/). It is noteworthy that AlphaFold2 predictions are not always as accurate as more traditional experimental methods and predict one stable conformation per protein, ignoring any dynamic changes in the protein structures. Therefore, the presented tripartite GT1b-Syt1-SV2 complex for selective binding and endocytic targeting of BoNT/A should be regarded as a predictive model only, and will have to be confirmed by other methods.

### QUANTIFICATION AND STATISTICAL ANALYSIS

Statistical tests were conducted in Prism 9 for macOS version 9.1.1. The normality of the data was tested with Kolmogorov-Smirnov tests unless otherwise stated, and non-parametric tests and parametric tests were used to compare two independent groups (Mann-Whitney test or t test) and multiple groups (Kruskal-Wallis test or ordinary one-way ANOVA multiple comparison test). All the presented super-resolution results in dot plots are from separate technical replications (*i.e.* from separate cultures) from the indicated number of independent experiments. Dot plots indicate average ± SEM from individual acquisitions or ROIs, unless otherwise stated. Additional details on sample sizes for each experiment can be found in the description of each experiment in the Methods section and respective figure legends.

**Image adjustments for figures and preparation of manuscript figures**

Brightness and contrast of acquired images was adjusted in ImageJ/Fiji and Adobe Photoshop 22.4.3 release (Adobe). Pseudo-coloring of multi-channel acquisition was done in ImageJ/Fiji. Figures were made in Adobe Illustrator 25.4.1 release (Adobe).

### Table S1: CRISPRi oligonucleotides

| Name | Sequence (5’->3’) | Description |
|---|---|---|
| Vec-pLV_R2 | TCCTTAATCAGCTCGCTCATAGGGCCGGGATTCTCCTCCAC | Reverse primer for dCAS9-KRAB linearization |
| Vec-pLV_F2 | ACTGGGGCACAAGCTTAATTGACCAGCACACTGGCGGC | Forward primer for dCAS9-KRAB linearization |
| TagBFP_F | CCTATGAGCGAGCTGATTAAGGAGAACATGC | Forward primer for TagBFP2 linearization |
| TagBFP_R | TCAATTAAGCTTGTGCCCCAGTTTGCTAG | Reverse primer for TagBFP2 linearization |
| RnSyt1-sgRNA1_F | CACCGCGTGCCTCGCACCGGTCCGCGG | Forward primer for gRNA1 against 5’-UTR of rat synaptotagmin-1 gene |
| RnSyt1-sgRNA1_R | AAACCCGCGGACCGGTGCGAGGCACGC | Reverse primer for gRNA1 against 5’-UTR of rat synaptotagmin-1 gene |
| RnSyt1-sgRNA2_F | CACCAGTACTCGCGTGCCTCGCACCGG | Forward primer for gRNA2 against 5’-UTR of rat synaptotagmin-1 gene |
| RnSyt1-sgRNA2_R | AAACCCGGTGCGAGGCACGCGAGTACT | Reverse primer for gRNA2 against 5’-UTR of rat synaptotagmin-1 gene |
| RnSyt1-sgRNA3_F | CACCTCCTCCTGCAGCGGCAGCATCGG | Forward primer for gRNA3 against 5’-UTR of rat synaptotagmin-1 gene |

| | | |
|---|---|---|
| RnSyt1-sgRNA3_R | AAACCCGATGCTGCCGCTGCAGGAGGA | Reverse primer for gRNA3 against 5'-UTR of rat synaptotagmin-1 gene |
| Lenti_F | CGTACCGGTTAGTAATGATCGACAATCAACC | Forward primer used to amplify the pLenti6.3 |
| Lenti_R | CGGAACTCCCAAGCTTATCGATAAAATTTTGA | Reverse primer used to amplify the pLenti6.3 |
| prom-Syn1_F | TTATCGATAAGCTTGGGAGTTCCGCTGCAGAGGGCCCTGCGTATGAG | Forward primer used to amplify synaptotagmin-1 with synapsin-1 promoter |
| RnSyt1_R | TGTCGATCATTACTAACCGGTACGTTACTTCTTGACAGCCAGCATGGCATCAAC | Reverse primer used to amplify synaptotagmin-1 with synapsin-1 promoter |

### Table S2: CRISPRi plasmids

| Unique identifier | Name | Description |
|---|---|---|
| AAAA-0238 | pLenti6.3-Syt1$^{wt}$-pH | *Rattus norvegicus* synaptotagmin1 under human synapsin1 promotor |
| AAAA-0240 | pLV-hUV6-sgRNA-dCas9-KRAB-Puro | Basic vector Addgene Cat. #71236 |
| AAAA-0244 | pLV-hUV6-sgRNA-dCas9-KRAB-TagBFP2 | Nontargeting control vector with TagBFP2, this paper |
| AAAA-0245 | pLV-RnSyt1-sgRNA1-dCas9-KRAB-TagBFP2 | Synaptotagmin-1 sgRNA1 with TagBFP2, this paper |
| AAAA-0246 | pLV-RnSyt1-sgRNA2-dCas9-KRAB-TagBFP2 | Synaptotagmin-1 sgRNA2 with TagBFP2, this paper |
| AAAA-0247 | pLV-RnSyt1-sgRNA3-dCas9-KRAB-TagBFP2 | Synaptotagmin-1 sgRNA3 with TagBFP2, this paper |

**Captions for Movies**

**Movie S1. Electron tomographic analysis of the hippocampal neurons following CRISPRi Syt1 KD and induced uptake of BoNT/Ai$^{wt}$-HRP.** Syt1 KD was performed using CRISPRi sgRNA1 Syt1 KD in cultured hippocampal neurons (on DIV14 for 7 days), after which neurons were stimulated for 5 min with high $K^+$ buffer supplemented with 5 µg $mL^{-1}$ BoNT/Ai$^{wt}$-HRP (dark precipitate), washed with low $K^+$ buffer and chased for 10 min. Neurons were then fixed, cytochemically stained and processed for EM. Approximately 200 nm thick resin sections was subjected to electron tomography. The section shows an axonal segment on the left, and a presynaptic area on the right. The plasma membrane is modeled in green, synaptic vesicles in cyan, microtubules in yellow and tubular structures containing BoNT/A$^{wt}$-HRP precipitate (highlighted in purple within yellow bounding boxes) in red. Bar 50 nm.

**Movie S2. Ribbon structure of BoNT/Ai.** Mapped lysine-residues (magenta spheres), E224A/R363A/Y366F light chain (LC) mutations (orange spheres; toxin inactivation), and heavy chain (HC) receptor binding mutations W1266L (BoNT/A$^{PSG}$) and G1141D/G1292R (BoNT/A$^{SV2}$) (cyan spheres).

**Movie S3. Dual-color single-molecule uPAINT imaging of BoNT/Ai$^{wt}$-At647N and SV2A$^{wt}$-pH/At565nb in live hippocampal neurons.** Hippocampal neurons expressing SV2A$^{wt}$-pH were stimulated with high $K^+$ buffer supplemented with 100 pM BoNT/Ai$^{wt}$-At647 and 100 pM anti-GFP At565nb and the mobility of SV2A-pH–bound At565nb (cyan) and the holotoxin (magenta) was recorded by TIRF microscopy (50 Hz, 20 ms exposure time). Bar 10 µm and playback 50 frames $s^{-1}$.

**Movie S4. Internalized BoNT/Ai$^{wt}$-At647N imaged with sdTIM in live hippocampal neurons** Hippocampal neurons were stimulated for 5 min with high $K^+$ buffer supplemented with 1 nM BoNT/Ai$^{wt}$-At647 (magenta), washed, chased for 10 min in low $K^+$ buffer, and imaged by TIRF microscopy (50 Hz, 20 ms exposure time). Playback 50 frames $s^{-1}$. Hippocampal neurons were transfected with EGFP to outline transfected neurons from the underlying cultured neurons and a representative image of the EGFP is superimposed on the acquisition (white).

**Movie S5. Retrograde transport of BoNT/Ai$^{wt}$-At647N in live hippocampal neurons.** Hippocampal neurons were stimulated with high $K^+$ buffer supplemented with 100 nM BoNT/Ai$^{wt}$-At647 for 5 min, washed in low $K^+$ buffer, chased for 2 h in medium and imaged with spinning-disk confocal microscopy. Retrograde (from nerve endings towards the soma; right to left) transport carriers positive for BoNT/Ai$^{wt}$-At647N are highlighted in magenta (circles), and the respective tracks are shown in multiple colors. Bar, 10 µm and playback 50 frames $s^{-1}$.

**Movie S6. AlphaFold prediction of tripartite GT1b-Syt1-SV2 nanocomplex and BoNT/A.** Predictive model of the assembled GT1b-Syt1-SV2A complex with bound BoNT/A on the plasma membrane. SV2A$^{T84A}$, Syt1$^{K326A,K328A}$ and Syt1$^{K52A}$, as well as the BoNT/A W1266L and G1141D/G1292R used in this study, are indicated.
